# Machine learning predictions of MHC-II specificities reveal alternative binding mode of class II epitopes

**DOI:** 10.1101/2022.06.26.497561

**Authors:** Julien Racle, Philippe Guillaume, Julien Schmidt, Justine Michaux, Amédé Larabi, Kelvin Lau, Marta A. S. Perez, Giancarlo Croce, Raphaël Genolet, George Coukos, Vincent Zoete, Florence Pojer, Michal Bassani-Sternberg, Alexandre Harari, David Gfeller

## Abstract

CD4^+^ T cells orchestrate the adaptive immune response against pathogens and cancer by recognizing epitopes presented on MHC-II molecules. The high polymorphism of MHC-II genes represents an important hurdle towards accurate prediction and identification of CD4^+^ T-cell epitopes in different individuals and different species. Here we collected and curated a dataset of 627,013 unique MHC-II ligands identified by mass spectrometry. This enabled us to precisely determine the binding motifs of 88 MHC-II alleles across human, mouse, cattle and chicken. Analysis of these binding specificities combined with X-ray crystallography refined our understanding of the molecular determinants of MHC-II motifs and revealed a widespread reverse binding mode in MHC-II ligands. We then developed a machine learning framework to accurately predict binding specificities and ligands of any MHC-II allele. This tool improves and expands predictions of CD4^+^ T-cell epitopes, and enabled us to discover and characterize several viral and bacterial epitopes following the aforementioned reverse binding mode.

## Introduction

CD4^+^ T cells are key components of the adaptive immune system. They are implicated in priming and modulating the natural immune response against pathogens and cancer. CD4^+^ T cells also play an essential role in cancer immunotherapy (Alspach et al., 2019; Borst et al., 2018), as demonstrated by CD4+ T-cell responses following neoantigen-based cancer vaccines (Hu et al., 2021; Ott et al., 2017; Sahin et al., 2017) and CD4^+^ T cell mediated regression of metastatic cancer following adoptive transfer of tumor-infiltrating lymphocytes (Tran et al., 2014; Zacharakis et al., 2018). CD4^+^ T cell activation starts with the recognition of epitopes presented by the highly polymorphic class II Major Histocompatibility Complex (MHC-II) on the surface of antigen presenting cells. Despite their central role in infectious diseases, autoimmunity and cancer, epitopes presented on MHC-II and targeted by CD4^+^ T cells are still poorly described and difficult to predict. This represents an important bottleneck for fundamental immunology, cancer immunotherapy and personalized cancer vaccines.

Peptides presented on MHC-II are processed by the class II antigen presentation pathway. Most of these peptides come from extracellular proteins ingested and degraded by the cell in the endocytic pathway (Neefjes et al., 2011). After cleavage, peptides typically 12-25 amino acids (AAs) long are loaded on MHC-II and the peptide-MHC-II complexes are displayed on the cell surface. The loading of peptides on MHC-II is facilitated by the action of chaperones, including HLA-DM and HLA-DO in human (Neefjes et al., 2011). The binding site of MHC-II molecules has been extensively characterized by X-ray crystallography and MHC-II ligands adopt a conserved binding mode in these structures. This canonical binding mode consists of a linear 9-mer binding core, which makes most of the interactions with the MHC-II binding site, and peptide flanking residues that extend on the N- and C-terminal parts of the binding core (Figure 1A) (Holland et al., 2013). Specific pockets are known to accommodate residues at anchor positions (mainly P1, P4, P6 and P9) in the binding core of the MHC-II ligands (Figure 1A) (Holland et al., 2013). Exceptions to this conserved binding mode have been reported in chicken where one MHC-II allele accommodates peptides with a 10-mer binding core (Halabi et al., 2021). In human, three peptides have been reported to bind in both the canonical and the reverse orientation (i.e. from N-to C-terminus and from C- to N-terminus) (Günther et al., 2010; Schlundt et al., 2012). It is however unclear how relevant and frequent this reverse binding mode is for naturally presented MHC-II ligands and CD4^+^ T-cell epitopes.

**Figure 1.**
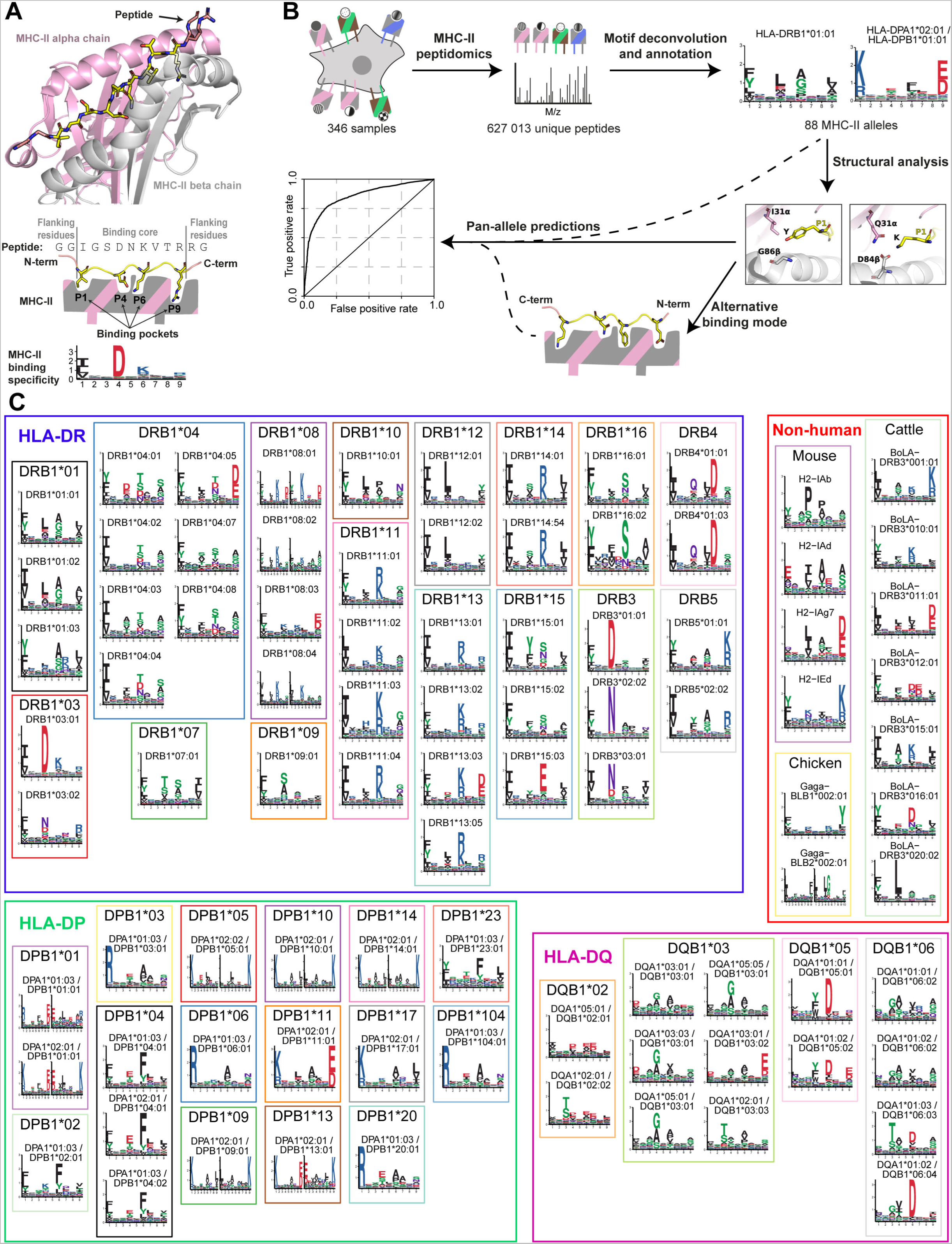
Curation of MHC-II peptidomics data reveals binding specificities for 88 MHC-II alleles. (A) Representative crystal structure of an MHC-II dimer (HLA-DRA*01:01 – DRB1*03:01) in complex with a peptide. The binding core of the peptide is shown in yellow, the peptide flanking residues in salmon, the MHC-II alpha chain in pink and the MHC-II beta chain in gray. Anchor positions P1, P4, P6, and P9 point towards the different MHC-II binding pockets and are visible in the binding motif below. (B) Schematic view of the MHC-II motif analysis and class II epitope prediction pipeline. The main steps of our analyses consist of (i) collection of a large dataset of naturally presented MHC-II ligands identified in multiple MHC-II peptidomics samples, (ii) motif deconvolution and annotation, (iii) structural analysis of MHC-II binding specificities and (iv) development of a machine learning predictor of MHC-II ligands and CD4+ T-cell epitopes. (C) Motifs describing the binding specificities of the 88 MHC-II alleles determined in this study. Multiple specificities, when identified for a given allele (e.g., HLA-DRB1*08:01), are shown side by side.

In human, the MHC-II is also called class II human leukocyte antigen (HLA-II) and consists of three gene loci directly involved in presenting antigens to CD4^+^ T cells: HLA-DR (including HLA-DRA1 and HLA-DRB1/3/4/5 genes), HLA-DP (including HLA-DPA1 and HLA-DPB1 genes) and HLA-DQ (including HLA-DQA1 and HLA-DQB1 genes). Except for HLA-DRA1, these genes are highly polymorphic and more than 9,100 alleles have been identified (IMGT/HLA database (Robinson et al., 2020) as of 23.06.2022). Within each gene loci, MHC-II form protein heterodimers composed of an alpha chain (e.g., HLA-DPA1*02:01) and a beta chain (e.g., HLA-DPB1*01:01). Due to the combinatorial between the alpha and beta chains, this leads to an even much higher number of possible heterodimers of MHC-II alleles (hereafter referred to as “MHC-II alleles” for simplicity) expressed on the cell surface in different individuals. The polymorphic residues mostly lie in the peptide binding site (Unanue et al., 2016), resulting in highly allele-specific peptide binding motifs (Abelin et al., 2019; Racle et al., 2019; Reynisson et al., 2020).

The polymorphism of MHC-II genes, the vast diversity of MHC-II binding motifs and the complexity of the class II antigen presentation pathway represent important hurdles for reliable predictions of naturally presented MHC-II ligands and CD4^+^ T-cell epitopes. In recent studies, we and others have shown how high-throughput mass spectrometry-based MHC-II peptidomics can be used to improve these predictions (Abelin et al., 2019; Chen et al., 2019; Racle et al., 2019; Reynisson et al., 2020). This was achieved through the identification of MHC-II motifs using either monoallelic samples (Abelin et al., 2019) or motif deconvolution in polyallelic samples (Racle et al., 2019; Reynisson et al., 2020). Beyond MHC-II binding motifs, MHC-II peptidomics also revealed specificity in the first and last AAs in peptide flanking residues of naturally presented MHC-II ligands, a specific peptide length distribution (peaked at 15 AAs) and a preference for a peptide binding core offset slightly shifted towards the C-terminus of the MHC-II ligands (Barra et al., 2018; Ciudad et al., 2017; Falk et al., 1994; Racle et al., 2019). The level of expression of both the epitope source proteins and the HLA-II molecules in antigen-presenting cells was also shown to correlate with antigen presentation (Abelin et al., 2019; Chen et al., 2019).

Today, several MHC-II ligand prediction tools are available. These include allele-specific (e.g., MixMHC2pred-1.2, limited to 38 alleles (Racle et al., 2019) and NeonMHC2, limited to 35 HLA-DR alleles (Abelin et al., 2019)), pan-HLA-DR (e.g., MARIA (Chen et al., 2019)) and pan-allele predictors (e.g., NetMHCIIpan-4.0 (Reynisson et al., 2020) and MHCnuggets (Shao et al., 2020)). The latter aim at capturing correlation patterns between MHC-II binding sites and binding specificities. However, the data used to train these predictors are still sparse, limited to few alleles and consists mostly of peptides presented by HLA-DR alleles. As a result, epitope predictions for poorly characterized alleles, especially HLA-DP and HLA-DQ alleles or alleles from other species, have limited accuracy.

Here, we collected and curated a very large dataset of MHC-II ligands and determined the binding specificities of more than eighty MHC-II alleles. Integrating these data with a machine learning framework enabled us to improve our molecular understanding and prediction capability of MHC-II ligands (Figure 1B). These results refine and expand our understanding of the universe of CD4^+^ T-cell epitopes that could be therapeutically targeted in infectious disease, autoimmunity and cancer immunotherapy.

## Results

### Curation of MHC-II peptidomics data reveals binding specificities for 88 MHC-II alleles

To improve our understanding of the specificity of MHC-II alleles and class II antigen presentation, we first performed a thorough literature curation to search for available mass spectrometry-based MHC-II peptidomics datasets and collected data coming from 30 published studies for a total of 322 samples and 615,361 unique peptides (Table S1, Data S1A). Most of these samples were obtained from human cells using anti-HLA-DR or anti-pan-HLA-II antibodies. Other samples were obtained using anti-HLA-DP or anti-HLA-DQ antibodies (Balen et al., 2020; Bergseng et al., 2015; Ritz et al., 2018), or cells transfected with tagged HLA-II allowing for the isolation of peptides bound to a single allele (Abelin et al., 2019). A few samples were obtained from mouse (Draheim et al., 2017; Sofron et al., 2016; Wan et al., 2020), cattle (Fisch et al., 2021) and chicken (Halabi et al., 2021). To further enrich for HLA-DP and HLA-DQ ligands, we used mass spectrometry-based MHC-II peptidomics to sequentially isolate peptides with anti-HLA-DR, anti-HLA-DP, anti-HLA-DQ and anti-pan-HLA-II antibodies (See Material and Methods). Applying this strategy to six different cell lines or meningioma tissues enabled us to obtain 44,334 unique peptides, including 11,779 HLA-DP and 16,146 HLA-DQ ligands (Table S1, Data S1B). This was especially useful with respect to the limited number of publicly available HLA-DQ ligands (31,045 unique peptides).

Combining all these data led to a total of 627,013 unique peptides (1,540,995 peptides when counting duplicates across samples) coming from 346 samples corresponding to 201 different cell lines or tissues, making it the largest currently available collection of mass spectrometry-based MHC-II peptidomics data (Data S1, Table S1). We then performed motif deconvolution with MoDec (Racle et al., 2019) on each of these samples (see Material and Methods). We could confidently describe the binding specificities of 88 MHC-II alleles, including 43 HLA-DR, 18 HLA-DP, 14 HLA-DQ, 4 mouse H-2, 7 cattle BoLA-DR and 2 chicken Gaga-BLB alleles (Figure 1C). Figure S1A shows that motifs identified in multiple samples sharing a common allele are highly reproducible. Our data also show that the distributions of peptide lengths and binding core offsets are conserved across both human and non-human MHC-II alleles (Figure S1B-C).

### MHC-II binding specificities reflect biochemical properties of the MHC-II binding pockets

Our large dataset of MHC-II binding motifs provides a unique opportunity to better understand the characteristics of MHC-II binding specificities. Consistent with previous studies, we observed that the HLA-DR, HLA-DP, mouse and cattle alleles usually have four clear anchors at positions P1, P4, P6 and P9. HLA-DQ have slightly weaker binding specificities with main anchors at P3, P4 and P6 and sub-anchors at P1 and P9, in general (Figure 1C). The binding specificities for all human, mouse and cattle alleles can be described by 9-mer motifs, confirming the binding core size of 9 AAs usually considered for MHC-II and observed in existing crystal structures. For the chicken allele Gaga-BLB2*002:01 that had been described earlier with a longer binding core (Halabi et al., 2021), we additionally searched with MoDec for a 10-mer motif. We observed that the binding specificity of this allele could be well described by two motifs, with almost half of ligands possessing a binding core of 9 AAs and the other ones having a binding core of 10 AAs (Figure S2A).

To investigate the molecular determinants of MHC-II binding specificities, we performed unsupervised clustering of the binding motifs for all human alleles for each HLA locus (i.e., HLA-DR, -DP and -DQ) and for each anchor position (Material and Methods). We could observe different classes of specificities (Figure 2A, left logos). We then retrieved the sequences of the most variable residues in the MHC-II binding pockets interacting with the anchor residue in the ligand for all alleles found in each cluster (Figure 2A, Table S2A-C, see Material and Methods). As expected, alleles found in distinct specificity clusters (i.e., distinct rows within each column of Figure 2A) also had differences in their binding pockets. To structurally interpret these different clusters, we used available crystal structures of representative alleles in each cluster (Figure 2A). For a few cases, no crystal structure was available, and we used structural modeling instead (see Material and Methods). This revealed a remarkable correspondence between the MHC-II binding motifs and the binding pockets of MHC-II alleles. For example, for HLA-DR alleles, large and bulky AAs (mainly F/Y) are observed at P1 when glycine (G) is found at 86β in the P1 binding pocket, while less bulky hydrophobic AAs (I/L/V) are observed at P1 in HLA-DR alleles when valine (V) is found at 86β in the binding pocket (Figure 2A, Table S2A). This mutual exclusivity reflects the steric clash that would happen between F/Y in the ligand and V in the binding pocket. For HLA-DP alleles, a small AA at 84β (G) correlates with bulky AA at P1 (F/L/Y/I), and a negatively charged AA at 84β (D) correlates with positively charged AA at P1 (K/R), with a clear salt bridge between the two opposite charges (Figure 2A, Table S2B). Furthermore, a long polar AA at 31α (Q) correlates with K at P1, while a long non-polar AA at 31α (M) correlates with R at P1 (Figure 2A, Table S2B). This can be explained by the fact that K at P1 can simultaneously engage into polar or charged interactions with D84β and Q31α, whereas the two nitrogens of R at P1 would preferentially face the two oxygens of the carboxyl group of D84β (Figure S2B). This conformation is more favorable if M is found at 31α instead of Q. Similar analyses between the MHC-II binding motifs at other anchor positions and the residues in the corresponding binding pockets are detailed in Table S2D. Remarkably, most of the observations in Figure 2A can be rationally explained in terms of steric hindrance or polar and charged interactions. These results demonstrate a clear correspondence between our MHC-II motifs and the sequences of MHC-II binding sites.

**Figure 2.**
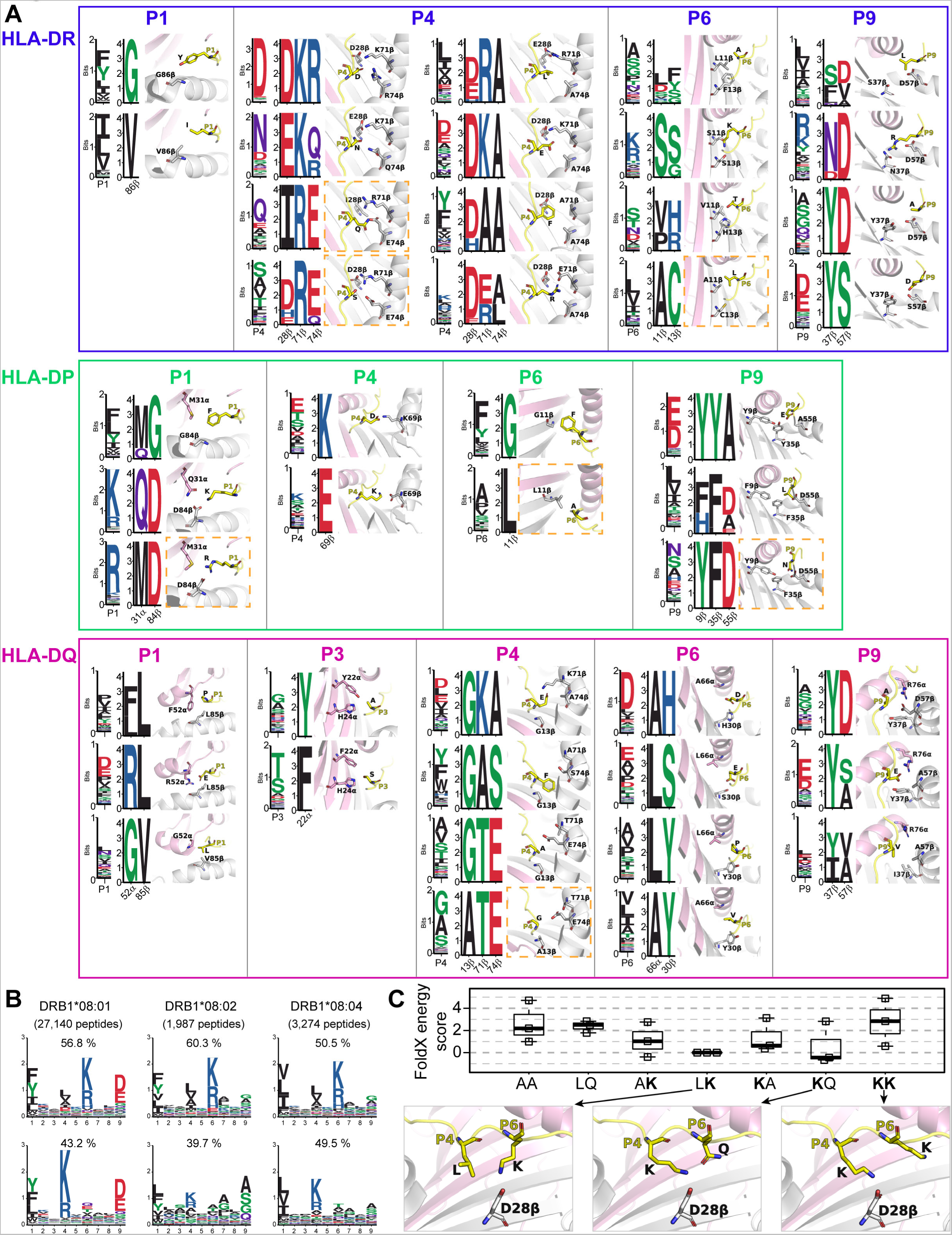
MHC-II binding specificities reflect biochemical properties of the MHC-II binding pockets. (A) For each type of human MHC-II (HLA-DR, -DP and -DQ) and each anchor position, MHC-II alleles were clustered based on the binding specificity at this position and the different clusters are shown as different rows for each position. On each row, the first motif (e.g., P1) represents the average peptide binding specificity of the MHC-II alleles in a cluster. The second motif (e.g., 86β) represents the sequence of the MHC-II residues making important contacts with the residue in the ligand at the specified anchor position. The structural arrangement of the residues in the MHC-II binding site (pink for alpha chain, grey for beta chain) and the residue in the ligand (yellow) is shown on the right based on existing X-ray structures. When no PDB structure was available, structural modeling was used instead and is indicated by a dashed orange rectangle. Details of the clusters of alleles at each position can be found in Table S2. (B) Multiple binding specificities for HLA-DRB1*08 alleles. Percentage above the motifs indicate the fraction of peptides assigned to each sub-specificity. (C) Molecular interpretation of the multiple specificities observed in HLA-DRB1*08:01. The top boxplot shows the calculated change in the FoldX energy score for various peptides with 0, 1 or 2 positively charged AAs at P4 and P6 (simulations based on three different peptides for each case, see Table S2E). The bottom part shows a model of HLA-DRB1*08:01 which contains one single negatively charged residue (D28β) able to interact with positively charged sidechains at either P4 or P6, but not both simultaneously.

For most alleles, a single binding specificity was observed. Yet, for several HLA-DRB1*08 we observed two binding motifs (Figure 2B). The two motifs suggest that a positively charged AA (K/R) is favorable either at P4 or P6, but not at both positions at the same time. To understand the molecular mechanism of this bi-specificity, we calculated the FoldX energy score of several variants of peptides with a charged residue either at P4 or P6, two charged residues at these positions or no charged residue (see Material and Methods). Consistent with the observed bi-specificity, our calculations indicate that peptides with two charged residues have higher energy scores (i.e., weaker predicted binding affinity) than those with only one charged residue (Figure 2C). This can be explained by the fact that the central part of the binding site of HLA-DRB1*08 alleles contains only one negatively charged residue (D28β) that can interact with positively charged sidechains either at P4 or P6, but not at both P4 and P6 (Figure 2C). This mutual exclusivity of charged residues at P4 and P6 appears to be further restricted to HLA-DRB1*08 alleles and may require G at 13β (Figure S2C-D).

### MHC-II binding specificities reveal a widespread reverse binding mode in MHC-II ligands

Another type of bi-specificity was observed for several HLA-DP alleles (Figure 1C). For these alleles, superimposing the motifs revealed a clear symmetry between the first and the second binding motifs (Figure 3A). It is unlikely that residues with opposite biochemical properties (e.g., K versus E at P1) could fit at the same position (e.g., P1 binding pocket). Therefore, we hypothesized that the second motif corresponds to a reverse binding mode where the peptides are bound from the C-terminus to the N-terminus. This hypothesis is further supported by the fact that the distribution of binding core offsets for peptides following the second motifs is skewed towards the N-terminus, unlike all other MHC-II ligands for which it is skewed towards the C-terminus (Figure S3A).

**Figure 3.**
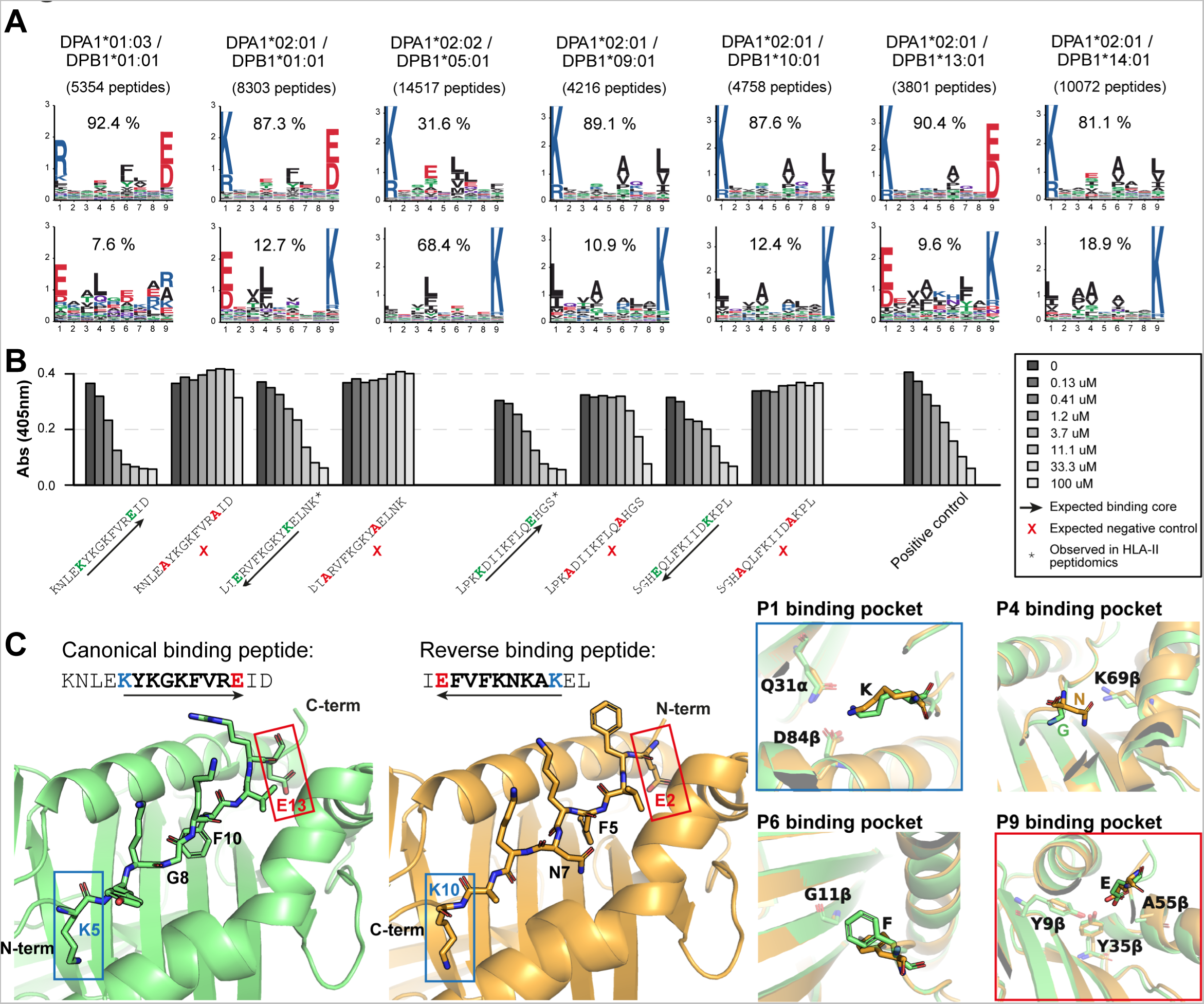
MHC-II binding specificities reveal a widespread reverse binding mode in MHC-II ligands. (A) HLA-DP alleles with symmetrical multiple binding specificities. Percentage above the motifs indicate the fraction of peptides assigned to each sub-specificity. (B) Binding competition assays of peptide variants bound to HLA-DPA1*02:01-DPB1*01:01. The different peptides correspond to (1) predicted canonical binders, (2) alanine mutations at anchor residues P1 and P9, (3) predicted reverse binders (reversed sequences) and (4) alanine mutations at P1 and P9 in the reversed sequence. Additional peptides are shown in Figure S3D. Stars indicate peptides that were seen in MHC-II peptidomics data. (C) Crystal structures resolved in this work of the canonical binder KNLE<u>KYKGKFVRE</u>ID and the reverse binder I<u>EFVFKNKAK</u>EL bound to HLA-DPA1*02:01-DPB1*01:01. The four panels on the right show the overlap of the two structures at the four binding pockets.

To validate our hypothesis, we first tested the binding of different peptides to HLA-DPA1*02:01-DPB1*01:01 (see Material and Methods). Starting with two peptides predicted to bind in the canonical orientation, we could validate their binding (Figure 3B). Mutation to alanine of the predicted P1 and P9 anchor residues abrogated the binding, confirming that these residues act as anchors, while replacing K at P1 with R did not affect the binding (Figure S3B). We then reversed the sequences, in order to obtain peptides following the second binding motif. Figure 3B shows that the binding was preserved. In general, the binding was stronger for peptides following the first motif (i.e., predicted canonical binders) than for those with the reversed sequence (i.e., predicted reversed binders). These results are fully consistent with the two motifs observed in MHC-II peptidomics data and their respective weights (Figure 3A). As a sidenote, one should not conclude that any ligand can be flipped and still bind, as demonstrated in Figure S3B with two peptides for which the binding was weaker.

We then attempted to crystallize peptides predicted to bind in either the canonical or the reverse orientation (Material and Methods). We first obtained the X-ray crystal structure (resolution 1.62 Å) of a peptide which matches the first motif (KNLE<u>KYKGKFVRE</u>ID, core underlined). This peptide binds in the canonical orientation to HLA-DPA1*02:01-DPB1*01:01, with K5 filling the P1 binding pocket and E13 filling the P9 binding pocket (Figure 3C-left, Figure S3C). We then crystallized a peptide compatible with the second motif (I<u>EFVFKNKAK</u>EL, resolution 2.9 Å). We observed that the binding happens in the reverse orientation with the first residue of the core (E2) filling the P9 binding pocket, and the last residue of the core (K10) filling the P1 binding pocket (Figure 3C, Figure S3C). The interactions mediated by each anchor residue were preserved, except for F in the P6 binding pocket that adopted a slightly different conformation in the reverse binder (Figure 3C). Overlaying the two structures further demonstrates a remarkable alignment not only of the sidechains (Figure S3D) but also of the backbone N-H and C=O groups (Figure S3E), as well as a conservation of most backbone H-bonds with the MHC-II binding site between canonical and reverse MHC-II ligands (Figure S3F). These results demonstrate that several HLA-DP alleles can accommodate peptides in both orientations and that these two binding modes are accurately captured with the bi-specificity observed in MHC-II ligands.

During class II antigen presentation, MHC-II ligands are processed by different proteases and footprints of this process are visible in the N- and C-terminal contexts of MHC-II ligands (Ciudad et al., 2017; Paul et al., 2018). However, the timing and the impact of the structural positioning of peptides in the MHC-II binding site are still unclear (Bird et al., 2009; Sercarz and Maverakis, 2003). MHC-II ligands binding in the reverse orientation provide an opportunity to shed light on this process. Figure S3G shows that the motifs of the N- and C-terminal contexts of the reverse binding peptides are very similar to those from the canonical binding peptides. Consistent with predictions from previous studies (Abelin et al., 2019), this observation supports a model where most of the cleavage and trimming takes place first, independent of the positioning of the peptides in the MHC-II binding site.

### MHC-II binding specificities can be accurately predicted for alleles without known ligands

Due to the high polymorphism of MHC-II genes, the binding specificity cannot be determined experimentally for all alleles. Our large and curated collection of MHC-II motifs provides an opportunity to capture correlation patterns between binding motifs and binding sites, and to train an accurate predictor of MHC-II ligands for any allele (referred to as pan-allele predictor). To this end, we designed a machine learning framework composed of two distinct successive blocks (Figure 4A). In the first block, the aim is to predict the MHC-II binding motifs (mathematically defined as Position Probability Matrices (PPMs), see Material and Methods) directly from the MHC-II sequences. In the second block, the aim is to predict actual MHC-II ligands based on their sequence and the PPM of the corresponding MHC-II allele.

**Figure 4.**
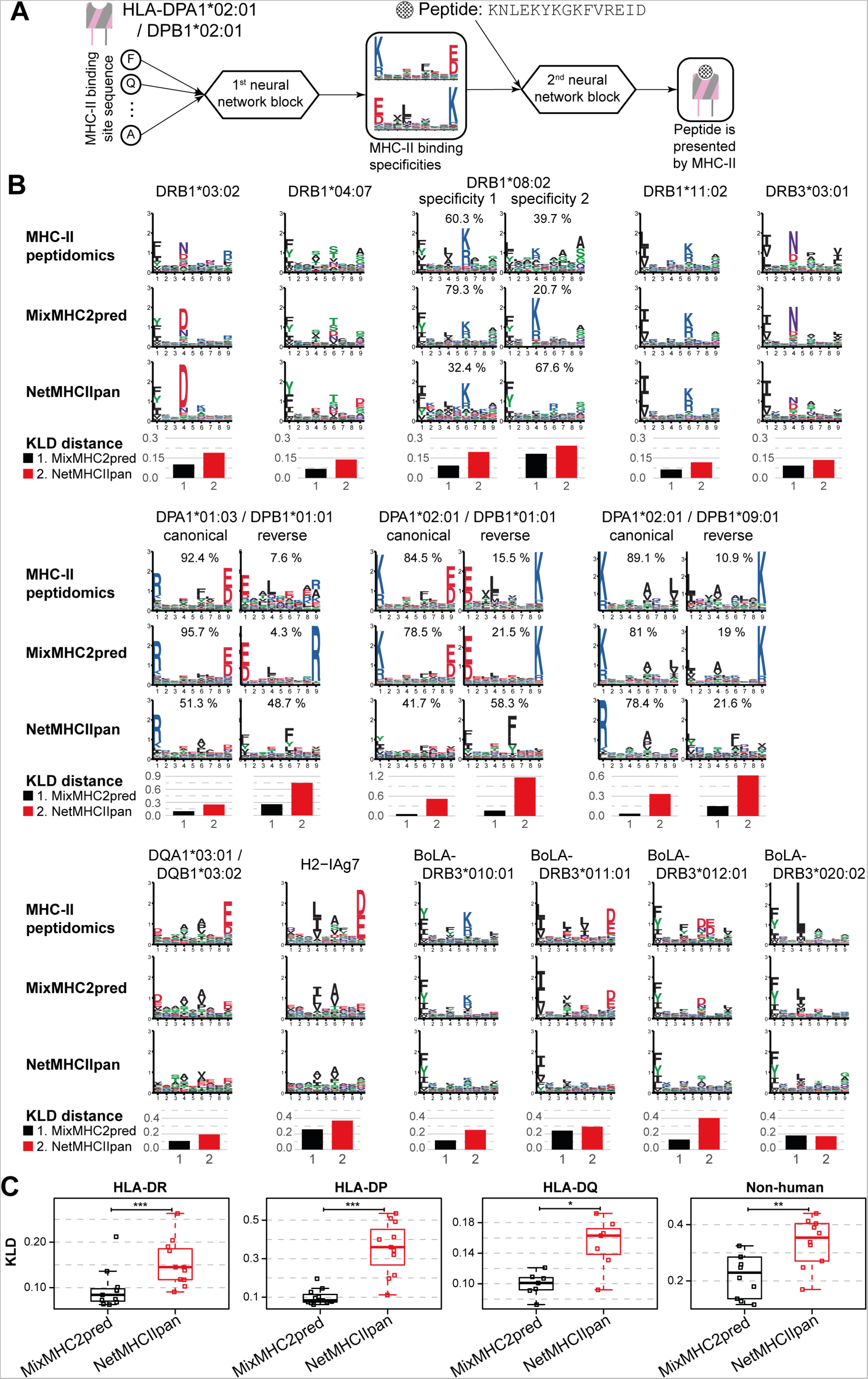
MHC-II binding specificities can be accurately predicted for alleles without known ligands. (A) Schematic description of the pan-allele predictor comprising two consecutive blocks of neural networks. In the first block, the input corresponds to the sequence of the binding site of the MHC-II allele, and the output corresponds to the PPMs describing the allele binding specificity. The second block combines the score of the peptide with the PPM of the MHC-II allele together with other features of MHC-II ligands (length, position of the core, N- and C-terminal contexts) to predict MHC-II presentation based on peptide sequences (see Fig. S4A for the full details of the model). (B) Comparison between actual and predicted motifs for alleles observed in monoallelic MHC-II peptidomics samples and absent from NetMHCIIpan training set (leave-one-allele-out cross-validation for MixMHC2pred). Multiple specificities, when present, are shown and the fraction of peptides observed and predicted per specificity is indicated above each motif (for NetMHCIIpan the multiple specificities were analyzed with MoDec, see Material and Methods). The average distance, measured with Kullback-Leibler divergence (KLD) per peptide core position, between the motifs observed in MHC-II peptidomics and the predicted ones is shown below each allele (see Figure S4B for similar analyses for additional alleles obtained from multiallelic samples). (C) KLD between the specificities observed in MHC-II peptidomics and predicted by MixMHC2pred (leave-one-allele-out) and NetMHCIIpan for all MHC-II alleles absent from NetMHCIIpan training. Box plots indicate the median, upper and lower quartiles; results of a paired two-sided Wilcoxon signed rank-test are indicated (*P<0.05, **P < 0.01, ***P < 0.001).

The first block consists of a set of fully connected neural networks for each binding core position separately (Figure S4A-left, see Material and Methods). The binding sites corresponding to each position and used as inputs of the neural networks were determined based on existing crystal structures (see Material and Methods). This block also incorporates the different multiple specificities and was trained on our set of PPMs representing each motif (see Material and Methods). The second block of the predictor consists of a fully connected neural network (Figure S4A-right). It takes as input the score of the peptide against the PPMs of the corresponding MHC-II allele together with other features linked to antigen presentation (i.e., peptide length, binding core offset and peptide processing/cleavage features, see Material and Methods). This block is trained on our dataset of MHC-II ligands (positives) and randomly selected peptides from the human proteome (negatives) (see Material and Methods).

As a first validation, we benchmarked how accurate our predictor, referred to as MixMHC2pred-2.0, was in predicting the MHC-II binding motifs for alleles without known ligands. To this end, we performed a leave-one-allele-out cross-validation, where all the data from one allele is removed from the training (see Material and Methods). To compare with the current state-of-the-art pan-allele predictor NetMHCIIpan-4.0, we focused on MHC-II alleles absent from its training set. For both predictors, predicted motifs were built by considering 100,000 random human peptides and selecting the top 1% best predicted peptides (see Material and Methods). The resulting motifs were compared with those derived from MHC-II peptidomics studies using the Kullback-Leibler divergence (KLD) (see Material and Methods). Our results show that MixMHC2pred better inferred the binding specificities of new alleles for both human and non-human MHC-II alleles (Figures 4B-C, S4B). Of note, the multiple specificities could be well predicted by MixMHC2pred, while they were not detectable in NetMHCIIpan predictions (Figures 4B, S4B, see Material and Methods).

### MixMHC2pred improves predictions of MHC-II ligands and CD4^+^ T-cell epitopes

We then benchmarked the accuracy of MixMHC2pred to predict naturally presented MHC-II ligands, using the leave-one-allele-out cross-validation setting (see Material and Methods). The Area Under the Receiver Operating Characteristic curve (ROC AUC) was computed as a measure of prediction accuracy. Results showed improved predictions for MixMHC2pred compared to other pan-allele predictors, both in human (Figure 5A-B) and non-human samples (Figure 5C). We next examined if MixMHC2pred would be amenable to predictions for species without known MHC-II ligands for any allele. To this end, we retrained our predictor removing all data coming from one species and predicted the MHC-II ligands from this species (referred to as leave-one-species-out cross-validation, see Material and Methods). We compared the predictions of this model with the predictions obtained in the leave-one-allele-out setting as well as to the full model, where all available data from all species were used in the training (Figure 5D). As expected, the full model was more accurate, followed by the leave-one-allele-out model and then the leave-one-species-out model. Nevertheless, AUC values were still much better than random when the predictor was not trained on any data from a given species, demonstrating that MHC-II ligand predictions can be extrapolated to new species, although with some loss in prediction accuracy (Figure 5D).

**Figure 5.**
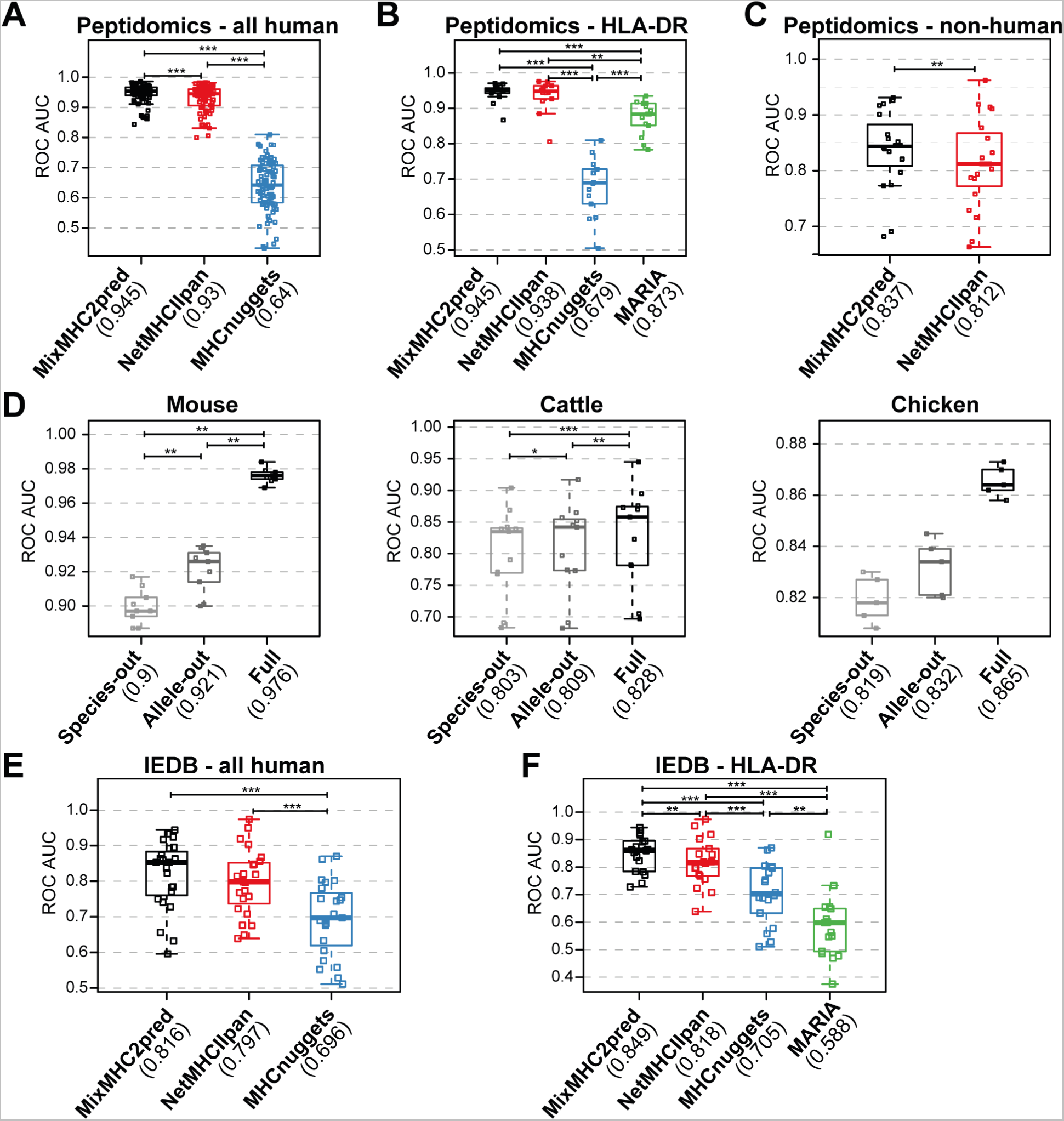
MixMHC2pred improves predictions of MHC-II ligands and CD4^+^ T-cell epitopes. (A-C) Accuracy (ROC AUC) for predictions of peptides presented by MHC-II. Only samples that are absent from the training set of all predictors are included. MixMHC2pred results were obtained in a leave-one-allele-out context. Results for (A) all human samples; (B) human HLA-DR only samples and (C) all non-human samples (mouse, cattle and chicken samples). MARIA can only be applied on HLA-DR and MHCnuggets only on human and mouse alleles. They are therefore included only in the corresponding analyses. (D) Accuracy for predictions of peptides presented by MHC-II in mice, cattle, and chickens, using our prediction framework trained on (i) all data except those from the species where predictions are made (leave-one-species-out), (ii) all data except those containing the allele for which predictions are made (leave-one-allele-out), or (iii) all data (full model). (E-F) Accuracy for predictions of CD4^+^ T-cell epitopes found in IEDB database, for (E) all human data, (F) only HLA-DR alleles. Numbers in parentheses below each predictor’s name correspond to the average ROC AUC values. Box plots indicate the median, upper and lower quartiles; the results of a paired two-sided Wilcoxon signed rank-test are indicated (*P<0.05, **P < 0.01, ***P < 0.001).

We then benchmarked the predictions for CD4^+^ T-cell epitopes (i.e., peptides presented by MHC-II and recognized by CD4^+^ T cells), using data coming from the Immune Epitope Database (IEDB) (Vita et al., 2019) and considering epitopes originating from viruses, bacteria and cancer that have been tested for CD4^+^ T cell immunogenicity. We computed the accuracy of the different predictors for each allele separately, based on the peptides that were annotated as immunogenic or not (see Material and Methods). Even though many of these epitopes had been selected based on exiting MHC-II ligand predictors (mainly NetMHCIIpan), we observed that the predictions of MixMHC2pred were more accurate than those of other tools (Figure 5E-F).

### Multiple specificities reveal reverse binding CD4^+^ T-cell epitopes

To capitalize on the ability of our tool to model multiple binding specificities of MHC-II alleles, we scanned common viral and bacterial proteomes and selected 39 peptides that were predicted to follow only the reverse binding mode of HLA-DPA1*02:01-DPB1*01:01 (Figure 6A, see Material and Methods). We stimulated CD4^+^ T cells isolated from PBMC of two HLA-DPA1*02:01-DPB1*01:01^+^ donors using pools of these peptides and measured cytokine production. After deconvolving the responses, we could identify seven peptides eliciting TNFα and IFNγ production (Figure 6B, Table S3A). We then built peptide-MHC-II multimers with four of those peptides. A clear multimer^+^ population was found for each epitope, demonstrating that the immunogenicity originated from the actual peptides bound to HLA-DPA1*02:01-DPB1*01:01 (Figure 6C, Material and Methods). To gain insights into the clonality of these reactive CD4^+^ T cells, we sequenced their T-cell receptor (TCR) alpha and beta chains. Oligoclonal responses were observed for each epitope (Table S3B), including a quasi-monoclonal recognition of the EBV epitope GELALTMRSKKLPIN (with a single TCRα and a dominating TCRβ). We next wondered if these TCR could be found in other donors. The alpha and beta chain sequences were used to query separately TCRα/TCRβ repertoires through the iReceptor web platform (Corrie et al., 2018) (see Material and Methods). Most of the alpha and beta CDR3 sequences of our study have been already observed in other donors, including 13 cases (out of 32) with exactly the same CDR3, V and J genes (Table S3B). Overall, these observations suggest that TCRs recognizing reverse binding epitopes are not rare in the human population.

**Figure 6.**
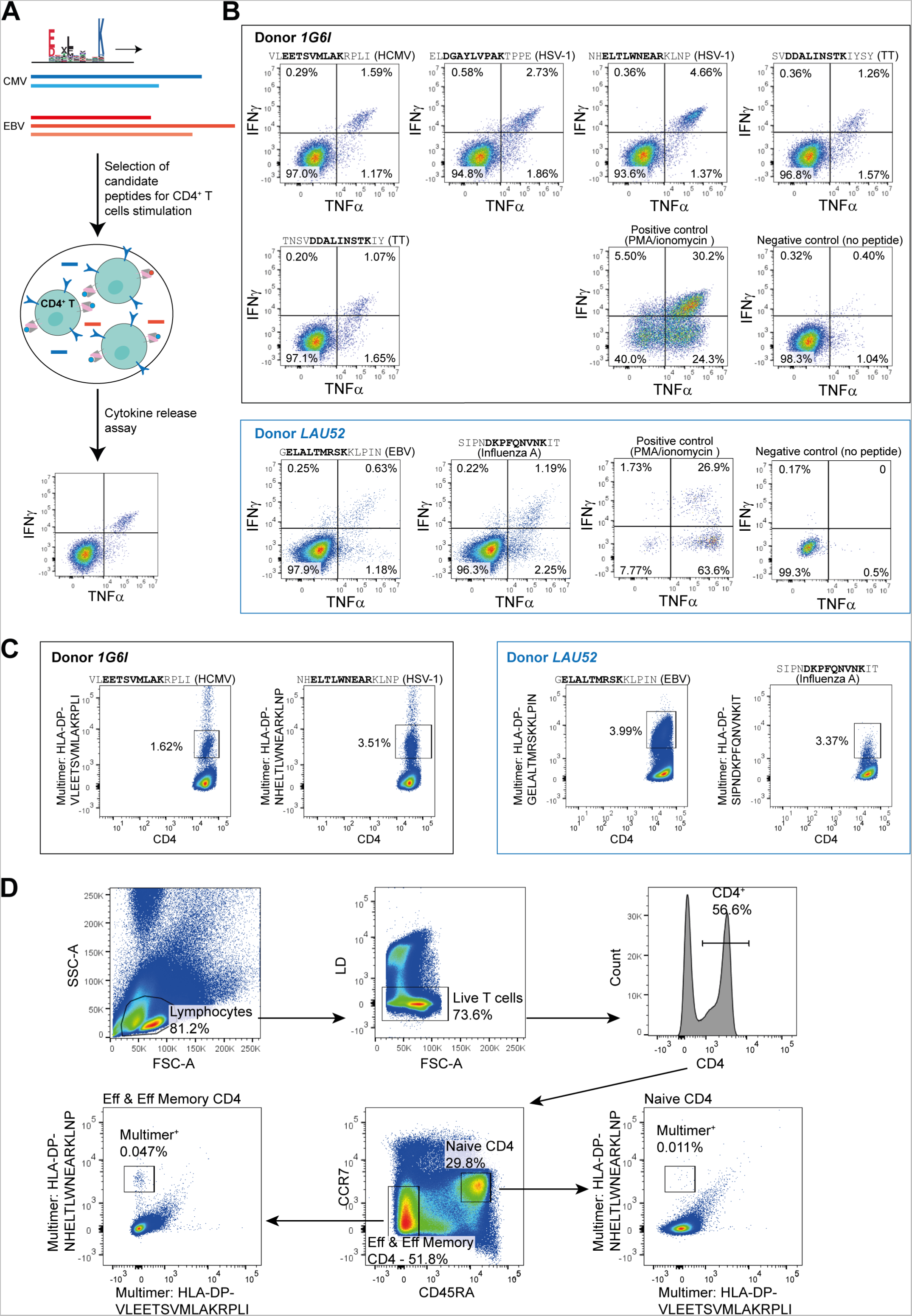
Multiple specificities reveal reverse binding CD4^+^ T-cell epitopes. (A) Schematics of the search strategy for reverse binding epitopes. Different viral and bacterial proteomes were scanned, searching for 15-mer peptides predicted to follow the reverse binding motif of HLA-DPA1*02:01-DPB1*01:01. CD4^+^ T cells from HLA-DPA1*02:01-DPB1*01:01^+^ donors were then stimulated with the selected peptides. Responses were evaluated through cytokine release assays deconvolving the response to the individual peptide. (B) TNFα and IFNγ response of the positive peptides observed after deconvolving the response in two HLA-DPA1*02:01-DPB1*01:01^+^ donors. The viral or bacterial species of origin of the peptide is indicated in parenthesis above each figure (EBV: Ebstein-Barr virus; HCMV: human cytomegalovirus; HSV-1: herpes simplex virus type 1; TT: tetanus toxin protein). (C) Validation with peptide-MHC-II multimers of the reverse binding epitopes (see Figure S5 for negative controls). (D) FACS results directly on *ex vivo* CD4^+^ T cells of donor *1G6I*, showing that recognition of the epitope NH<u>ELTLWNEAR</u>KLNP happens through effector and effector memory CD4^+^ T cells, not naive CD4^+^ T cells.

Next, we investigated whether these epitopes had elicited a memory response. To this end, we tested if the CD4^+^ T cells from the two donors could recognize these epitopes directly *ex vivo*, without any prior stimulation, using the four multimers validated above. For one donor we could observe a direct *ex vivo* response that was mediated by effector and effector memory CD4^+^ T cells (Figure 6D). This shows that reverse binding MHC-II ligands can elicit natural CD4^+^ T-cell recognition. Reverse binding epitopes identified in this work had very poor scores (%Rank > 20) with other MHC-II ligand predictors (Table S3A). This demonstrates how thorough analysis of large datasets of MHC-II ligands together with machine learning algorithms can improve and expand the scope of CD4^+^ T-cell epitope predictions.

## Discussion

CD4^+^ T cells play a central role in immune recognition of infected or malignant cells, and response to cancer immunotherapy. CD4**^+^** T cells and MHC-II alleles have also been linked with multiple autoimmune diseases, including rheumatoid arthritis, celiac disease, type 1 diabetes, multiple sclerosis and narcolepsy (Latorre et al., 2018; Matzaraki et al., 2017). However, identifying class II (neo-)epitopes displayed on MHC-II molecules and targeted by CD4**^+^** T cells remains challenging. This is especially true for epitopes displayed on HLA-DP and HLA-DQ alleles.

Here we capitalized on both public and in house MHC-II peptidomics data to derive accurate MHC-II motifs for a large panel of MHC-II alleles, including HLA-DP and HLA-DQ alleles whose binding specificity was less well defined in previous studies. The fact that the different classes of binding specificities at each anchor position could be rationalized in terms of molecular interactions with residues in the different binding pockets provides an independent validation of the MHC-II motifs derived directly from MHC-II peptidomics data. We also note that the HLA-DP ligands corresponding to reverse binders were not included in the analyses of Figure 2A, since our results subsequently revealed that AAs at P1, resp. P4, actually fit in the P9, resp. P6, binding pocket, and vice versa. Some of the correlation patterns between the MHC-II binding motifs and MHC-II binding pockets (Figure 2A) can already be found in the definition of MHC-II supertypes (Greenbaum et al., 2011). However, several other observations are specific to this work. For instance, HLA-DPB1*01:01 and HLA-DPB1*04:02 were assigned to the same supertype (Greenbaum et al., 2011), while both their specificity at P1 (K/R, resp. F/L/Y/I) and their P1 binding pockets (D, resp. G at 84β) are clearly different. This mainly reflects the limited number (<30 alleles) and lower resolution of MHC-II motifs used to define MHC-II supertypes. Of note, these limitations are also affecting state-of-the-art MHC-II ligand predictors (see examples in Figures 4B, S4B and S6). As such our results provide both a refined view of the main MHC-II binding specificities, in line with the recent classification proposed for HLA-DP alleles (Laghmouchi et al., 2022; Meurer et al., 2021), and a robust implementation of these observations into an accurate MHC-II ligand prediction tool (MixMHC2pred-2.0).

We observed that multiple specificities were especially frequent in HLA-DP alleles and we anticipate that a few multiple specificities may have been missed with our motif deconvolution approach for MHC-II alleles with less ligands (e.g., HLA-DPA1*02:01-DPB1*11:01, see also HLA-DRB1*08:03). These observations are consistent with the previously reported multiple specificity of HLA-DPA1*02:02-DPB1*05:01 (Balen et al., 2020). For HLA-DP alleles, our work demonstrates that the two different motifs correspond to two different binding modes (i.e., canonical and reverse) of HLA-II ligands. We do not exclude that reverse binders may also be found among the ligands of other alleles, similar to the CLIP peptides observed to bind in both orientations to HLA-DRB1*01:01 (Günther et al., 2010). However, we could never detect multiple motifs with the same type of symmetry for HLA-DR or HLA-DQ alleles. Moreover, most alleles do not have the same specificity at P1 and P9, or P4 and P6, suggesting that peptides fitting HLA-DR or HLA-DQ motifs mainly bind in the canonical orientation and that reverse binders, if present, comprise only a small fraction of the actual ligands. This potential paucity of the reverse binding mode in HLA-DR (and HLA-DQ) ligands could explain why the earlier observations of reverse binders based on some very specific peptides (Günther et al., 2010; Schlundt et al., 2012) have not been followed by other similar observations. Furthermore, this reverse binding mode was not captured by existing predictors and was not particularly anticipated from a structural point of view, since several contacts between MHC-II and their ligands are mediated by the backbone atoms of the ligands (Figure S3F), and would not necessarily be conserved in the reverse orientation. This demonstrates the power of unbiased analyses of MHC-II ligands to unravel novel properties of MHC-II alleles.

Within human, our work shows that MHC-II binding motifs and MHC-II ligands can be accurately predicted even for alleles without known ligands (Figures 4B-C, 5A-B, and S4B). This is an important improvement of MixMHC2pred-2.0 compared to the previous version, MixMHC2pred-1.2, as well as NeonMHC2, which are allele-specific predictors and can only be run for a small subset of alleles (24 HLA-DR, 10 HLA-DP, 4 HLA-DQ for MixMHC2pred-1.2, and 35 HLA-DR for NeonMHC2). As a result, these allele-specific predictors are not applicable to most patients and could not be included in our benchmarks. When attempting to make predictions in species without any information about MHC-II ligands (i.e., leave-one-species-out), we observed lower, though not random, accuracy. This is likely a limitation of all pan-allele MHC-II ligand predictors, which should be used with care in distant species like fish or birds. For instance, most tools would not be able to predict the 10-mer binding core of the chicken allele Gaga-BLB2*002:01, if not explicitly trained on these data.

Altogether, our work provides a unique high-quality dataset of MHC-II ligands, precise definition of MHC-II binding motifs, refined understanding of the molecular determinants of these motifs including a widespread reverse binding mode of HLA-DP ligands, and improved machine learning predictions of CD4^+^ T-cell epitopes. The fact that the viral epitopes following the reverse binding mode and eliciting responses in effector-memory CD4^+^ T-cells could not have been identified with other MHC-II ligand predictors demonstrates the promise of machine learning algorithms like MixMHC2pred to better characterize and expand the repertoire of CD4^+^ T-cell epitopes. The improved accuracy of CD4^+^ T-cell epitope predictions may contribute to accelerating personalized immunotherapy approaches in autoimmune diseases or cancer.

## Acknowledgments

We thank Camilla Jandus from the University of Geneva for providing us a patient-derived PBMC sample. We thank Simon Eggenschwiler from the University of Lausanne for help with the implementation of MixMHC2pred webserver. We thank the Protein Modeling Unit of the University of Lausanne for the support in Structural Bioinformatics. We thank Peter A. van Veelen, Michel G.D. Kester and their colleagues (Balen et al., 2020), Ana Marcu and her colleagues (Marcu et al., 2021), Birkir Reynisson, Morten Nielsen and their colleagues (Fisch et al., 2021), for sharing MHC-II peptidomics data relating to their published studies.

D.G. and JR acknowledge support from the Swiss Cancer Research Foundation (KFS-4104-02-2017). G.Croce is supported by the Marie-Curie fellowship (H2020-MSCA-IF-2020, No 101027973).

## Author contributions

Conceptualization, J.R. and D.G.; Methodology, J.R.; Software, J.R.; Investigation, J.R., P.G., J.S., J.M., A.L., K.L., M.A.S.P., G.Croce, F.P., and D.G.; Resources, G.Coukos, V.Z., F.P., M.B.-S., A.H., and D.G.; Data Curation, J.R.; Writing – Original Draft, J.R. and D.G.; Writing – Review & Editing, J.R. and D.G.; Visualization, J.R., J.S., and D.G.; Supervision, J.R., G.Coukos, V.Z., F.P., M.B.-S., A.H., and D.G.; Funding Acquisition, D.G.

## Declaration of interests

The authors declare no competing interests.

## Material and Methods

### Curation of publicly available MHC-II peptidomics data

We searched in the literature and on ProteomeXchange (http://www.proteomexchange.org/) (Deutsch et al., 2020) for available high-throughput mass spectrometry-based MHC-II peptidomics datasets in which the MHC-II typing was also known.

We downloaded the sequences of different reference proteomes from EMBL-EBI (https://ftp.ebi.ac.uk/pub/databases/reference_proteomes/): human (UP000005640_9606), mouse (UP000000589_10090), cattle (UP000009136_9913) and chicken (UP000000539_9031), obtaining both the canonical and additional sequences in fasta format (from release 2020_04). The peptides obtained from MHC-II peptidomics datasets were then mapped to these proteomes, in order to identify their protein of origin and determine their N-/C-terminal contexts (3 residues upstream of the peptide + 3 N-terminal residues of the peptide (N-terminal context); 3 C-terminal residues of the peptide + 3 residues downstream of the peptide (C-terminal context)).

### Cells and patient material

Epstein–Barr-virus-transformed human B-cell lines JY (ATCC, 77442), CM467, RA957, (a gift from P. Romero (Ludwig Institute for Cancer Research Lausanne), were maintained in RPMI-1640+GlutaMAX medium (Life Technologies) supplemented with 10% heat-inactivated FBS (Dominique Dutscher) and 1% penicillin–streptomycin solution (BioConcept). Cells were grown to the required cell numbers, collected by centrifugation at 1,200 rpm for 5min, washed twice with ice cold PBS and stored as dry cell pellets at *−*20 °C until use. PBMCs from donor 1G6I, PBMCs from melanoma patient LAU52 and snap-frozen meningioma tissues from patients (3865-DM, 3947-GA, 4021) were obtained from the bio-bank of the Centre Hospitalier Universitaire Vaudois (CHUV, Lausanne, Switzerland). Informed consent of the participants was obtained following requirements of the Institutional Review Board (Ethics Commission, CHUV). Protocol F-25/99 has been approved by the local ethics committee and the biobank of the Lab of Brain Tumor Biology and Genetics. Protocol 2017-00305 for antigen and T cell discovery in tumors has been approved by the local ethics committee. Protocol F-42/92 has been approved by the local ethics committee.

### HLA typing

Genomic DNA was extracted using DNeasy kit from Qiagen and 500ng of genomic DNA was used for the typing. High-resolution 4-digit HLA typing was performed with the TruSight HLA v2 Sequencing Panel kit from Illumina according to the manufacturer instruction. Briefly, class I and class II genes were amplified by PCR. Illumina adapters were added by tagmentation. After normalization and purification, the samples were sequenced on a MiniSeq instrument (Illumina). Sequencing data were analyzed with the Assign TruSight HLA v.2.1 software (Illumina).

### Generation of antibody-crosslinked beads

Anti-pan-HLA-II and anti-HLA-DR monoclonal antibodies were purified from the supernatant of HB145 (ATCC, HB-145) and HB298 cells (ATCC, HB-298), respectively, grown in CELLLine CL-1000 flasks (Sigma-Aldrich) using protein A-sepharose 4B beads (pro-A beads; Invitrogen) while anti-HLA-DP (Leinco Technologies) and anti-HLA-DQ (from either Biorad or MyBioSource) were purchased from the respective providers. Antibodies were cross-linked separately to pro-A beads at a concentration of 1 to 2 mg of antibodies per milliliter of beads following incubation with pro-A beads for 1h at room temperature. Chemical crosslinking was performed by addition of dimethyl pimelimidate dihydrochloride (Sigma-Aldrich) in 0.2M sodium borate buffer, pH 9 (Sigma-Aldrich) at a final concentration of 20mM for 30min. The reaction was quenched by incubation with 0.2M ethanolamine, pH 8 (Sigma-Aldrich) for 2h. Crosslinked antibodies were kept at 4 °C in PBS until use.

### Purification of HLA-II, HLA-DR, HLA-DP, HLA-DQ bound peptides

Cells were lysed in PBS containing 0.25% sodium deoxycholate (Sigma-Aldrich), 0.2mM iodoacetamide (Sigma-Aldrich), 1mM EDTA, 1:200 protease inhibitors cocktail (SigmaAldrich), 1mM phenylmethylsulfonylfluoride (Roche) and 1% octyl-beta-dglucopyranoside (Sigma-Aldrich) at 4 °C for 1h. The lysis buffer was added to cells at a concentration of 1×10^8^ cells per milliliter. Cell lysates were cleared by centrifugation with a table-top centrifuge (Eppendorf Centrifuge) at 4 °C at 14,200 rpm for 50min. Meningioma tissues were placed in tubes containing the same lysis buffer and homogenized on ice in three to five short intervals of 5 s each using an Ultra Turrax homogenizer (IKA) at maximum speed. For 1 g of tissue, 10–12ml of lysis buffer was required. Cell lysis was performed at 4 °C for 1h. Tissue lysates were cleared by centrifugation at 20,000 rpm in a high-speed centrifuge (Beckman Coulter, JSS15314) at 4 °C for 50min.

Tissue cleared lysates were loaded first on affinity purification columns (BioRad, 731-1550) containing pro-A beads (pre-clear column, to remove non-specific antibodies). Tissues and cells lysates were loaded sequentially on columns containing cross-linked beads in the following order: CM467 samples on anti-DQ, DP, DR, pan-HLA-II antibodies, 3865 and JY on anti-DR, DQ, DP, pan-HLA-II antibodies, and samples RA957, 3947-GA, 4021 on anti-DR,DP,DQ, pan-HLA-II antibodies. The affinity columns were then washed with 2 column volumes of 150 mM sodium chloride (NaCl; Carlo-Erba) in 20 mM Tris-HCl pH 8, 2 column volumes of 400 mM NaCl in 20 mM Tris-HCl pH 8, and again 2 column volumes of 150 mM sodium chloride in 20 mM Tris-HCl pH 8. Finally, the beads were washed in 1 column volume of 20 mM Tris-HCl pH 8. HLA complexes and the bound peptides were eluted by adding twice 1% trifluoroacetic acid (TFA) or 4 times Acetic acid 0.1N at a volume equivalent to or slightly higher than the volume of beads present in the column. HLA peptides were purified and concentrated with by loading into Sep-Pak tC18 96-well plates (Waters) preconditioned with 1 mL of 80% acetonitrile (ACN) in 0.1% TFA and then with 2 mL of 0.1% TFA. The C18 wells were then washed with 2 mL of 0.1% TFA. The HLA peptides were eluted with 500 *μ*L of 32% ACN in 0.1% TFA into Eppendorf tubes. Recovered peptides were dried using vacuum centrifugation (Thermo Fisher Scientific) and stored at *−*20°C.

### LC–MS/MS analyses of HLA-II peptides

HLA-II, HLA-DR, HLA-DP and HLA-DQ peptide samples were resuspended in 10 µl of 2% ACN in 0.1% formic acid (FA) and aliquots of 3 µl were used for each MS analysis. The LC-MS/MS system consisted of an Easy-nLC 1200 (Thermo Fisher Scientific) coupled to a Q Exactive HF-X mass spectrometer (Thermo Fisher Scientific). Peptides were separated on a 450-mm analytical column (8-*μ*m tip, 75-*μ*m inner diameter, PicoTip emitter, New Objective) packed with ReproSil-Pur C18 (1.9-*μ*m particles, 120 Å pore size, Dr Maisch GmbH). The separation was performed at a flow rate of 250 nl/min by a gradient of 0.1% formic acid in 80% acetonitrile (solvent B) and 0.1% formic acid in water (solvent A).HLA-II peptides were eluted by the following gradient: 0 to 80 min (2 - 32% B); 80 to 84 min (32 – 45% B); 84 to 85 min (45 – 100% B); and 85 to 95 min (100% B). Data were acquired using a data-dependent acquisition (DDA) method. Full-scan MS spectra were acquired in the Orbitrap at a resolution of 60,000 (at 200 *m/z*) with an auto gain control (AGC) target value of 3×10^6^ ions. For Tandem mass spectrometry (MS/MS), ten most abundant precursor ions were sequentially isolated, activated by higher-energy collisional dissociation (NCE = 27) and accumulated to an AGC target value of 2×10^5^ with a maximum injection time of 120 ms. In the case of assigned precursor ion charge states of one, and from six and above, no fragmentation was performed. MS/MS resolution was set to 15,000 (at 200 *m/z*). Selected ions were dynamically excluded for additional fragmentation for 20 s. The peptide match option was disabled. The raw files and MaxQuant output tables will be deposited to the ProteomeXchange Consortium via the PRIDE (Perez-Riverol et al., 2022) partner repository.

### Peptide identification

We employed the MaxQuant platform v.1.5.5.1 (Cox and Mann, 2008) to search the peak lists against a fasta file containing the human proteome (Homo_sapiens_ UP000005640_9606, the reviewed part of UniProt, including 21,026 entries downloaded in March 2017) and a list of 247 frequently observed contaminants. Peptides with a length between 8 and 25 amino acids were allowed. The second peptide identification option in Andromeda was enabled. The enzyme specificity was set as unspecific. A false-discovery rate of 1% was required for peptides and no protein false-discovery rate was set. The initial allowed mass deviation of the precursor ion was set to 6ppm and the maximum fragment mass deviation was set to 20ppm. Methionine oxidation and N-terminal acetylation were set as variable modifications.

### Deconvolution and annotation of MHC-II motifs

To search for motifs describing the binding specificities of the alleles present in our compiled MHC-II peptidomics dataset (Table S1), we used our motif deconvolution tool MoDec (Racle et al., 2019). MoDec uses a probabilistic framework to search for common motifs of size L (L=9 in general for MHC-II ligands) present anywhere along the sequence of the MHC-II ligands identified by mass spectrometry in a given sample. The different motifs identified per sample typically correspond to binding specificities of the different MHC-II alleles. Both the mapping of each peptide to a specific motif and the identification of the binding core are derived from the maximum responsibility value returned by MoDec. Potential contaminant peptides are also identified during the deconvolution. MoDec-1.1 was run using the recommended options “--pepLmin 12 -- specInits --makeReport -r 50”, searching for motifs of length 9 AAs, and searching between 1 and up to 12 different motifs per sample (depending on the sample). For the chicken samples, where a bulging binding mode of 10 AAs binding core had been previously observed (Halabi et al., 2021), we additionally searched for 10-mers motifs.

Following our previously established procedure (Racle et al., 2019), all samples were manually reviewed and MHC-II motifs were annotated to their respective MHC-II allele by identifying shared motifs across samples sharing the same MHC-II allele. In this way, we could identify the motifs of 88 different alleles. Some samples were of too low quality to be included in the analyses (e.g., too few peptides present, or incorrect MHC-II typing where clear motifs are present but are describing other alleles than the alleles supposed to be present in the given sample), and these samples were not included in Table S1 nor Data S1. Further, peptides clearly assigned to motifs corresponding to alleles not supposed to be in the sample were considered as contaminants and were removed from further analyses. Peptides that could not confidently be assigned to a specific allele were not considered for the analyses of binding specificities. These include cases with too few peptides to build a clear motif, peptides assigned to the flat motif of MoDec, or when two alleles of similar binding specificity are present in a sample but a unique motif describing these alleles was obtained. These peptides were nevertheless kept for the prediction benchmarks of MHC-II ligands to prevent any potential biases in our validation datasets. Multiple specificities appear when two (or more) clearly different motifs from a sample are identified as coming from the same allele (either because the sample is monoallelic or because the same multiple specificity motifs are appearing in multiple samples sharing only the given allele). Motifs identified for each allele in each sample are shown in Figure S1A. Motifs shown in Figure 1C were obtained by grouping together the peptides assigned to each allele from all samples and aligning them based on the binding core identified by MoDec. ggseqlogo (Wagih, 2017) was used to plot these motifs in all the figures (with the height of the motifs corresponding to Shannon entropy measured in bits). The list of all peptides assigned to each allele in each sample can be found in Data S1.

### Motifs clustering

For each allele for which we could obtain a motif, we started by computing a position probability matrix, PPM*^a^*, (*a*: the allele, *l*: the binding core position, *i*: the amino acid identity), using all the identified binding cores of all the peptides assigned to this allele (Data S1). We further included a pseudocount based on the BLOSUM62 substitution matrix with a parameter *β*=200 (Nielsen et al., 2004). The Kullback-Leibler divergence (KLD) was then computed between all pairs of alleles from a same gene locus (HLA-DR, HLA-DP or HLA-DQ), for each binding core position *l*:

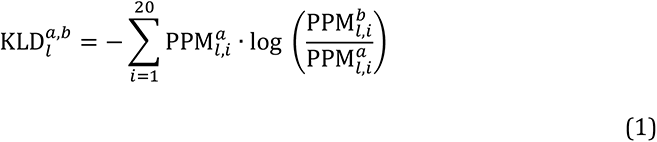

These KLD were then clustered through hierarchical clustering (using *hclust* function from R with the *average* clustering method). Thresholds to define the different clusters based on the hierarchical clustering were manually defined. Resulting clusters are given in Table S2A-C. The binding specificities plotted in Figure 2A (first motifs to the left in each column) correspond to the average PPM*^a^* between the alleles of each cluster. Clusters containing a single allele were not considered in the analysis.

### MHC-II sequences retrieval and alignment

Human MHC-II sequences were retrieved from IPD-IMGT/HLA database (Robinson et al., 2020).

Mouse MHC-II sequences (called H2-IAx and H2-IEx, where *x* gives the name of the allele) were manually retrieved from NCBI’s Protein database (https://www.ncbi.nlm.nih.gov/protein/), searching for MHC class II sequences of Mus musculus.

Cattle MHC-II sequences (BoLA-DR) were retrieved from IPD-MHC database (Maccari et al., 2017).

Chicken MHC-II sequences (Gaga-BL) are not yet part of IPD-MHC database. We found the accession numbers corresponding to the different alleles in the *Online Resource 2* of (Afrache et al., 2020), searched for these accession on NCBI’s Nucleotide database (https://www.ncbi.nlm.nih.gov/nuccore) and obtained the translated CDS sequences of the BLB1 and BLB2 genes.

The sequences were aligned together using the function *msaClustalOmega* from the *msa* package in *R* (Bodenhofer et al., 2015; Sievers et al., 2011).

### Analysis of available MHC-II structures, determining MHC-II-peptide contact positions

Crystal structures in PDB format where obtained from the RCSB PDB (https://www.rcsb.org/) (Berman et al., 2000; Rose et al., 2017). Structures containing human MHC-II alleles were retrieved based on the following *Sequence Motif*: “WRLEEFGRFASFEAQGALANIAVDKANLEIMTKRSNYTPITN” for HLA-DR, “FYVDLDKKETVWHLEEFG” for HLA-DP and “CLVDNIFPPVVNIT” for HLA-DQ. A custom script was used to determine to which MHC-II alpha and beta chains each PDB file corresponded (based on the chain A and chain B sequences in the PDB files and the MHC-II sequences obtained above).

We then manually reviewed these structures to determine the peptide binding cores. Residues in the MHC-II alpha and beta chains that were in close contact to each peptide binding core position (distance < 5 Å) were determined and kept in our list of contact positions if the same residue passed the distance threshold for at least two different MHC-II alleles and if the AA at this residue was not conserved among the MHC-II alleles for which we had obtained a binding motif. These residues in MHC-II alleles are those most likely to influence the binding specificity at a given binding core position in MHC-II ligands.

Sequence logos of the most important allele contact positions in each cluster (see above) were drawn with ggseqlogo (Wagih, 2017) (Figure 2A; Table S2A-C includes all contact positions). The numbering of the contact position residues follows the numbering found in X-ray structures for the alpha and beta chains. For the alleles from the first cluster of HLA-DQ at P1, we manually renumbered the amino acids F/L found at the residue 51α to exchange them with the gap found at 52α as they structurally better align to the residues at position 52α from the other HLA-DQ alleles.

We used PyMOL (https://pymol.org) to show representative images of the MHC-II and peptide contacts (Figures 1A-B, 2A), obtained from the following PDB IDs: 1BX2, 1DLH, 1KLU, 1S9V, 1UVQ, 2NNA, 3C5J, 3LQZ, 3PL6, 3WEX, 4IS6, 4MAY, 4MDJ, 4OZF, 4P57, 4P5M, 6BIR, 6CPL, 6DIG, 6PX6 and 7N19.

### Structural modeling

HLA-DR alpha chain, HLA-DR beta chain and peptide sequences were used as starting points for the modeling. Homology models of the HLA-II-peptide complexes were generated using Modeller software v10.1 (Webb and Sali, 2016). Template structures were retrieved from Protein Data Bank (Rose et al., 2017). Top matching templates were identified from the template library using an internal database with annotated alleles. The closest template was determined using the BLOSUM62 scoring function (Henikoff and Henikoff, 1992). A total of 2,000 models were produced for each HLA-II-peptide complex. These models were subsequently ranked based on the sum of the Discrete Optimized Potential Energy (DOPE) calculated using Modeller over the peptide residues, as well as the HLA residues within 6 Å from the peptide. For each HLA-II-peptide complex, molecular interactions were analyzed in the top 5 ranked models over the 2,000. The final HLA-II-peptide structural model corresponds to the one with best score and highest number of favorable interactions.

The effect of amino acid mutations on HLA-II-peptide structural stability was estimated using FoldX software after modeling the mutation using its buildmodel function (Schymkowitz et al., 2005). Changes in FoldX energy score (DG mutant – DG wild-type) were calculated for each mutant using an ensemble of 10 conformations. A change > 1kcal/mol means a destabilizing effect while a change < -1 kcal/mol means a stabilizing effect. In Figures 2C and S2C, differences in FoldX energy scores are relative to the “LK” case.

### Production of HLA-DPA1*02:01-DPB1*01:01

The extracellular region of the HLA-DPA1*02:01 and HLA-DPB101:01 chains were separately cloned into pMT\BiP\V5-His A (ThermoFisher scientific). The alpha chain construct harbors the acidic leucine zipper and terminates by a 6His-tag. The beta chain construct contains the basic leucine zipper and terminates with AviTag sequence. To generate cell lines expressing HLA-DPA1*02:01-DPB1*01:01, the two plasmids with a third plasmid conferring puromycin resistance, were cotransfected into Drosophila S2 cells using Cellfectin (ThermoFisher Scientific) according to the manufacturer protocol. Protein expression was induced by addition of 1 mM CuSO4. MHC class II molecules were purified from supernatants with Chelating Sepharose FF (Merck). Peptide loading was performed in citrate saline buffer (100 mM citrate, pH 6.0, 0.2% *β*-octyl-glucopyranoside (Calbiochem), 1×complete protease inhibitors (Roche)) with 100 µM peptide at 37°C for 24 hours, buffer-exchanged on a HiPrep 26/10 desalting column (Merck) into AviTag buffer and subsequently biotinylated with the BirA enzyme according to the manufacturer instructions (Avidity, Denver, Colorado, USA). Biotinylated MHC class II-peptide complexes were purified on a HisTrap HP column (Merck) and kept at -80°C until multimerization with streptavidin conjugates.

For protein crystallization “empty” HLA-DPA1*02:01-DPB1*01:01 was purified on a Superdex 75 10/300 GL column (Merck) into 20mM Tris pH 8.0, 150mM NaCl and concentrated at 10mg/ml.

### Binding competition assays

Four peptides following well one of the two observed binding specificities of HLA-DPA1*02:01-DPB1*01:01 and present in multiple MHC-II peptidomics samples containing this MHC-II allele were selected. These sequences were additionally “inverted” to have the same sequences but in the other orientation, in order to have four peptide sequences fitting well the observed canonical binding specificity and four sequences fitting well the reversed binding specificity. In addition, we also added the same sequences but where the AA present in the predicted P1 binding pocket was replaced by arginine (or lysine if the WT sequence had arginine), and we also added expected negative binders where the predicted peptide binding anchors P1 and P9 were replaced by alanine.

All these peptides were chemically synthesized using standard fmoc chemistry, purified by RP-HPLC (>80 % purity) and analyzed by UPLC-MS. Peptides were kept lyophilized at -80°C. To test the binding of these peptides to HLA-DPA1*02:01-DPB1*01:01, competition assays were performed by mixing in v-bottom 96-well plate (Greiner Bio-One) in 50 ml of citrate saline buffer (described above) 1 μg of the biotinylated empty allele with a FLAG-tagged peptide (IKTEKKTVQFSDDVQ) at fixed concentration of 2 μM and candidate peptides were added to each well to a final concentration of 0, 0.13, 0.41, 1.2, 3.7, 11.1, 33.3, and 100 μM. For the control, untagged peptide (IKTEKKTVQFSDDVQ) was used at the respective concentrations to the mix of allele and FLAG-tagged peptide. After incubation at 37 °C overnight. The binding of the tagged peptides to HLA-II molecule was measured by ELISA. The mix was transferred to a plate coated with avidin and the FLAG-peptide was detected with an anti-FLAG-alkaline phosphatase conjugate (Merck), developed with pNPP SigmaFAST (Merck) substrate and absorbance was read with a 405-nm filter.

### Protein crystallization, data collection, structure determination and model refinement

HLA-DP (HLA-DPA1*02:01-DPB1*01:01) protein at around 10 mg/ml was mixed with peptides at a final concentration of 10 mM, and co-crystallized by hanging drop vapor diffusion method. Crystals of HLA-DP-canonical binder (sequence: KNLEKYKGKFVREID) formed in a couple of weeks in 15% w/v PEG 4000, 0.2 M Magnesium chloride hexahydrate, 0.1M Sodium cacodylate pH 6.5 and crystals of HLA-DP-reverse binder (sequence: IEFVFKNKAKEL) in 8% w/v PEG 20K, 8% v/v PEG 500 MME, 0.2M Potassium thiocyanate, 0.1M Sodium acetate pH 5.5. The crystals were cryoprotected with 25% glycerol. Diffraction data were collected at the Paul Scherrer Institute (SLS, Villigen) at PXIII beamline. Data were processed with the XDS Program Package (Kabsch, 2010). Structures were solved by molecular replacement using Phaser-MR and PDB entry 3WEX as template model. Manual model building and structure refinement were carried out in Phenix Suite (Liebschner et al., 2019) using coot software (Emsley and Cowtan, 2004) and phenix-refine, respectively. After validation, the models were deposited in the PDB database. Data collection and refinement statistics are summarized (Table S4). The structures were displayed with PyMOL (https://pymol.org). The polar and charged interactions between the peptide and residues in the MHC-II binding site (Figures S2B, S3F) were determined with PyMOL, using default parameters.

### Motifs of the N- and C-terminal contexts of the HLA-DP ligands

Only peptides that were annotated as coming from an allele with the observed reversed binding mode were considered for this analysis (i.e., corresponding to the ligands shown in Fig. 3A). ggseqlogo (Wagih, 2017) was then applied on the N-terminal and C-terminal context sequences of these peptides, separately for the peptides following the canonical or reverse orientation (Figure S3G).

### Development of the pan-allele MHC-II ligand predictor

Our pan-allele predictor, MixMHC2pred-2.0 is composed of 2 successive blocks of neural networks with distinct tasks (Figures 4A, S4A). We implemented these neural networks in R, using the packages *keras* (version 2.7) and *tfdatasets* (version 2.7), relying on TensorFlow (version 2.6).

The first block describes the binding specificity. It consists of independent neural networks for each peptide binding core position, *l* (hereafter we refer to these independent neural networks as *NN^1^_l_*). The input of *NN^1^_l_* is the sequence of the contact AA residues in the P*_l_* binding pocket of the MHC-II allele *a*, and the output is the PPM at this binding core position for this allele (i.e. PPM*^a^*, including the BLOSUM62 pseudocount, described above). The contact AA residues used as input are the joint set of close contact residues between HLA-DR/-DP/-DQ explained above (Table S2A-C), after renumbering these based on the alignment of all the MHC-II sequences together, including human and non-human MHC-II. Each input AA was encoded to a numerical vector of size 21 equal to the sum of the one-hot encoding of the given AA plus the row corresponding to this AA in the BLOSUM62 probability matrix (Henikoff and Henikoff, 1992); the 21^st^ element of this vector represents a gap/absent AA (this 21^st^ element has value 0 except when the given “residue” in the allele is a gap instead of a true AA, in which case this last element has a value of 2). Each *NN^1^_l_* is composed of one fully connected hidden layer with 100 hidden nodes, based on rectified linear unit (ReLU) activation function and a gaussian noise of std 0.1. A dropout of 0.2 was added after these hidden nodes during the model training, and a softmax activation function was used for the output layer. The loss optimized corresponded to the Kullback Leibler divergence, and it was optimized using Adam optimizer with a learning rate of 0.005, a decay 0.005/250 and other parameters at default values. A maximum of 250 epochs was set and optimization stopped after no improvement of the loss during 30 epochs.

In order to account for the multiple specificities that are observed for some alleles, we replicated these *NN^1^_l_* three times: a first time including only all the canonical binders of specificity 1, a second time where the canonical binders of specificity 2 are used instead when the given allele had two canonical binders specificity (i.e. for the DRB1*08 alleles at present, but could accommodate additional if observed), and a third time where the reverse binders are used instead of the corresponding canonical binders when a reverse binding specificity was observed for the given allele (the sequence of the reverse binding peptides was inverted there, to have the correct peptides’ AA in the P1 binding pocket of the allele for example). A last independent neural network of the 1^st^ block is implemented to predict the fraction of peptides in these different binding specificities. The input here is the full list of contact residues from the MHC-II alleles, encoded in the same way as above and the output is the fraction of peptides observed in each of the 3 sub-specificities (canonical 1, canonical 2 or reverse) for the given allele. This neural network has similar structure than above’s *NN^1^_l_* except that 50 hidden nodes are used, with a learning rate of 0.0025, a maximum of 500 epochs and the loss corresponds to the categorical cross entropy. To avoid cases where multiple specificities would be present but with too few ligands to be observed, we restricted the training of this part to MHC-II alleles with more than 3,000 observed ligands. When predicting allele specificities, an MHC-II allele is assumed to possess multiple specificities only if this last neural network predicts a fraction of canonical specificity 2 or reverse specificity of at least 1% for the given allele. The training of all these neural networks of the 1^st^ block is repeated 5 times with same parameters and the final outputs are the average between these.

After having trained the first block, we can give the sequence of any MHC-II allele as input and this first block will return the predicted binding specificities of this allele (PPM*^a^*^,*s*^) (with *s* for the specificity: canonical 1, canonical 2 and reverse), as well as the relative fraction of peptides that are predicted to be bound to this allele in each of the three specificities (*w_s_*). The second neural network block, *NN^2^*, combines these with other features directly linked to a given peptide sequence (its sequence, length, binding core offset, peptide’s N-/C-terminal contexts), in order to predict if the peptide is presented by the given allele (Figure S4A - right). First, a PPM-based binding score is determined based on the given MHC-II allele specificities and peptide sequence:

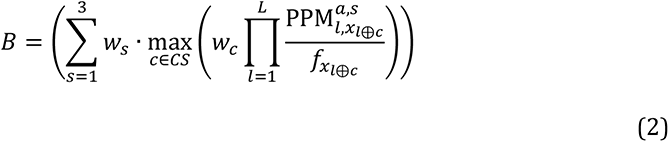

Where *w_c_* is the relative weight of the binding core offset *c* (similar to Figure S1C) and the best (maximum) value among all potential peptide binding core offsets is used for the inner parenthesis; *L* is the binding core length (i.e. 9 AAs); *x_j_* indicates which amino acid is found in the peptide at the position *j*; *f_i_* is a normalization factor, equal to the frequency of amino acid *i* in the human proteome. The “*l*⊕*c*” in *x_l_*_⊕*c*_ is the “special sum” previously defined (Racle et al., 2019), which makes that the binding core offsets are symmetric around 0 for each peptide (i.e., a binding core offset is equal to 0 when the binding core of the peptide is perfectly at the middle of the peptide, a negative value means the binding core is towards the peptide N-terminus and a positive value means the binding core is towards the peptide C-terminus). The binding score *B* of the peptide is then transformed to a percentile rank *B_rank_* based on the scores of 10,000 random human peptides of the same size. The 1^st^ input of *NN^2^* corresponds to a min-max scaling between 0 and 1 of the *log(B_rank_ + 10^-4^)* (where the min *B_rank_* is 0 and the max is 100, and 10^-4^ avoids *log(0)*). The 2^nd^ input of *NN^2^* is a one-hot encoding of the best binding core offset *c* (determined from Eq. (2) above), with values considered between -6 and 6. The peptide length is also one-hot encoded, for sizes between 12 and 21. The last set of inputs corresponds to the 12 AAs of the N- and C-terminal contexts, which were encoded following the same procedure as described above for the *NN^1^_l_* (in case of an unknown AA (“X”), the value 1/20 is used for the corresponding elements of this input vector).

Following these encoded input features, the *NN^2^* consists of a fully connected neural network with 1 hidden layer of 200 hidden nodes following a ReLU activation function with a gaussian noise of std 0.1. A dropout of 0.2 was added after these hidden nodes during the model training, and a sigmoid activation function was used for the single output node (with a value 1 if the given input peptide is presented by the given MHC-II alleles and 0 if not). Adam optimizer was used, with a learning rate of 0.001, a decay of 0.001/150 and other parameters at default values. The binary cross entropy loss was optimized. A validation split of 1/5 was used and early stopping after 50 epochs without validation loss improvement was set (or a maximum of 150 epochs otherwise).

To train this *NN^2^*, we used as positives the peptides observed in the MHC-II peptidomics samples (Data S1). We did not consider samples with missing MHC-II typing (e.g. samples obtained through anti-pan-HLA-II peptidomics when the HLA-DP had not been typed, but if the sample was obtained through anti-HLA-DR it was sufficient to have the HLA-DR typing), nor samples from chicken or cattle origin or containing a high fraction of predicted contaminant peptides, nor samples obtained through MAPTAC experiments (Abelin et al., 2019) (we observed some biases in their peptide length distributions towards longer peptides than expected). Only peptides of sizes 12-21 AAs long were kept, and peptides whose context could not be determined were also removed. We then downsampled the training set to keep a maximum of 200,000 positive peptides. In multiallelic samples, all potential MHC-II alleles were kept (i.e. we did not use the allele annotation from MoDec): equation (2) was applied to each allele of this sample and the best *B_rank_* score was used for the inputs of *NN^2^*. We finally removed peptides with best *B_rank_* > 30 (better binders have lower values), which likely correspond to contaminant peptides observed in MHC-II peptidomics. For negative peptides we used four times more random human peptides than positives, with a uniform length distribution between 12 and 21 AAs.

After its training, the scores of *NN^2^* are transformed to percentile ranks (%Rank) based on the scores of 10^6^ random human peptides and making that these follow the peptide length distribution observed in MHC-II peptidomics. For our final model, MixMHC2pred-2.0, the training of *NN^2^* is repeated 5 times and results correspond to the average of these repetitions. When running MixMHC2pred-2.0 in multiallelic samples, the %Rank against each allele is returned and the score of this sample is taken as the best (lowest) %Rank.

### Benchmarking of MHC-II binding specificity predictions

To compare the MHC-II binding specificities predicted by MixMHC2pred and NetMHCIIpan, we used 100,000 random human peptides of size 12-21 AAs and scored these using the different predictors against each allele of interest. For each allele, the best scoring 1% peptides were considered as the ligands to this allele, and their binding cores (returned by the predictor) were used to determine the frequency of each amino acid at each binding core position. We then compared these frequencies with the frequencies observed in MHC-II peptidomics data of the given allele by computing the KLD for each peptide binding core position between these frequencies (Eq. (1)). These KLD were averaged between all binding core positions. Lower KLD values mean that the predicted frequencies are closer to the frequencies observed in MHC-II peptidomics data. In Figure 4B, the comparison is performed against MHC-II peptidomics data coming only from monoallelic samples, while in Figures 4C and S4B it includes all MHC-II peptidomics data based on above’s annotation of the motifs after the deconvolution with MoDec. MARIA and MHCnuggets were not included in these analyses as their output only consists of the predicted peptide presentation score, but they do not predict the binding core, which prevents inferring the motifs. Likewise, MixMHC2pred-1.2 and NeonMHC2 were not included in these analysis as they are allele-specific predictors and therefore cannot do any prediction for an allele that would be left-out from their training set in a leave-one-allele-out context.

To determine how accurate these specificities are for new MHC-II alleles, only alleles that were absent from NetMHCIIpan training were considered here. In this respect, we trained MixMHC2pred in a stringent leave-one-allele-out cross-validation setting: when doing the predictions for allele *A*, no peptide annotated as coming from allele *A* is used during the training of the first predictor block *NN^1^*, and all peptides coming from all samples containing this allele *A* are removed from the second predictor block *NN^2^* (i.e., even if the peptide is annotated as coming from another allele in this sample, as long as the allele *A* is part of the list of alleles from this sample).

Multiple specificities were considered, and the fraction of peptides observed in each sub-specificity (when present) in MHC-II peptidomics data is indicated above the corresponding motifs (Figures 4B, S4B). One of the outputs of MixMHC2pred tells the predicted sub-specificity for each peptide and we directly used this. NetMHCIIpan does not return any information about from which sub-specificity a peptide would be coming. In order to allow having multiple specificities for NetMHCIIpan as well, we applied MoDec on the binding cores of the top 1% best predicted peptides from NetMHCIIpan, (running MoDec with the options “--nruns 50 --makeReport --specInits --no_flat_mot”). For alleles possessing multiple or reverse specificities, we show in Figure 4B and S4B the two motifs determined in this way for NetMHCIIpan, while for the alleles possessing a single specificity the motifs are directly obtained from NetMHCIIpan without applying MoDec. In Figure S4B, we show two rows of results for NetMHCIIpan, corresponding to using directly its results (i.e., with a single motif) or applying MoDec on its binding cores as described here.

### Benchmarking of MHC-II ligand predictions

The benchmark in Figure 5A-C was performed using the data from the various MHC-II peptidomics samples (Table S1, Data S1). All peptides from a given sample were used together with the set of alleles describing this sample. Peptides of sizes 12-21 were considered. We did not include samples with missing MHC-II typing, the *AUT01_xx* samples from (Marcu et al., 2021) due to high predicted contamination, nor samples from MAPTAC experiments (Abelin et al., 2019) due to observed biases described above. The positives were the peptides observed in each sample and we added four times more random peptides as negatives, taken from the same proteins as the proteins observed in the positive peptides and following a uniform length distribution between 12 and 21 AAs. In multiallelic samples, the scores of all these peptides for all alleles were computed and the best score among the sample’s alleles was kept (lowest %Rank for MixMHC2pred, lowest %Rank_EL for NetMHCIIpan, lowest ic50 for MHCnuggets and highest score for MARIA). Using the predicted scores of each peptide, the area under the curve of the Receiver Operating Characteristic curve (ROC AUC) was computed for each predictor and for each sample separately.

To avoid using the same peptides in testing and training of the predictors, we considered only samples that were absent from NetMHCIIpan and MARIA’s training sets (MHCnuggets is not trained on any of these samples as it only considers binding affinity data). In this way, the tested samples were therefore absent from the training of NetMHCIIpan, MARIA and MHCnuggets, but the MHC-II alleles from these test samples were often still part of the training of these predictors (i.e., the same alleles were present in some other samples used in the training of these predictors, and therefore the specificity of these alleles could already be well described by these predictors). For our predictor we used the same stringent leave-one-allele-out cross-validation setting than described above, where the %Rank of each allele was obtained separately based on this leave-one-allele-out setting and then the best %Rank among those was used, ensuring that no peptide coming from any sample containing the test allele was present in its training set.

For MARIA, the gene expression of each protein from which a peptide is originating is needed as further input. The gene expression pre-defined in MARIA were used, based on our annotation of from which type of tissue each sample/cell line is originating (BRCA, COAD, K562, …).

To compare the prediction accuracy for new species (Figure 5D), the test sets were composed of all MHC-II peptidomics samples from the given species (mouse, cattle or chicken) and random negative peptides, as described above. In addition to the leave-one-allele-out model and to the full model (trained on all peptides from all samples described above), we also trained leave-one-species-out models, where all data from the given test species were removed from the training and the model was trained on the remaining data only.

### Benchmarking of CD4^+^ T-cell epitope predictions

All data for human CD4+ T cells from the IEDB database were downloaded (as of 06.08.2021). We then filtered this data to keep the peptides of sizes 12-21 AAs which were observed in multimer/tetramer, ICS and ELISPOT assays, and whose 4-digits MHC-II typing had been determined, considering the “Allele evidence codes”: “MHC binding assay”, “Single allele present”, “T cell assay - Mismatched MHC molecules/ Biological process measured/ MHC subset identification/ T cell subset identification” and “Statistically inferred by motif or alleles present”. This dataset included directly peptides annotated either as positives or negatives, and no artificial negatives were added for this analysis. The ROC AUC was computed for predictions made per allele separately, keeping only alleles with at least 2 positive and 2 negative peptides. As in the experiments from this dataset the short peptides were usually directly tested, the antigen presenting cells did not need to cleave these peptides before presentation. Therefore, the part relating to context encoding is not meaningful and we used here the option of not encoding this peptide context in NetMHCIIpan. MixMHC2pred includes a similar option, which consists in internally replacing the AAs from the context by “X”s and where the %Ranks are recomputed accordingly.

### Searching for candidate CD4^+^ T-cell epitopes following the reverse binding mode

We downloaded viral proteomes from UniProt (The UniProt Consortium, 2021) (https://www.uniprot.org/), from EBV (Ebstein-Barr virus), HCMV (human cytomegalovirus), HSV-1 (herpes simplex virus type 1), HSV-2 (herpes simplex virus type 2), Influenza A virus (only from the HA and NA proteins), SARS2 (SARS-CoV-2) and VZV (Varicella-zoster virus), considering only the reviewed proteins, potentially coming from multiple strains of the given viruses. We also downloaded from UniProt the tetX gene of tetanus toxin protein (TT), which is produced by the bacteria clostridium tetani.

These proteomes were then cut in all overlapping 15-mer peptides, and we computed the binding scores of all these peptides with HLA-DPA1*02:01-DPB1*01:01, keeping separate the scores from the canonical and reverse specificity (i.e., scores from Eq. (2) but keeping the scores from the index *s* separate instead of summing over them, corresponding to the canonical and reverse specificity). We then selected a set of 4-5 peptides per proteome that had a good score towards the reverse specificity and only a weak score towards the canonical specificity and we synthesized these peptides for experimental testing.

### Peptides and peptide-MHC-II multimers

Peptides and peptide-MHC-II multimers were produced by the Peptide & Tetramer Core Facility of the University Hospital of Lausanne (CHUV). Peptides were chemically synthesized using standard fmoc chemistry, purified by RP-HPLC (>90 % purity) and analyzed by UPLC-MS. Peptides were kept lyophilized at -80°C. Biotinylated peptide-MHC-II monomers, loaded with peptides of interest were multimerized using streptavidin-PE (Cat# SA10044, Thermofisher Scientific) or streptavidin-APC (Cat# 405207, Biolegend) conjugates, then stored at 4°C and used within a week.

### Identification of antigen-specific CD4^+^ T-cell responses

CD4^+^ T cells were isolated (ref 130-045-101, Miltenyi) from cryopreserved PBMC and co-incubated (10^6^ mL^-1^) with autologous irradiated CD4-depleted PBMCs (10^6^ mL^-1^) and pools of 3 to 4 peptides (2 µM) in RPMI supplemented with 8 % human serum and IL-2 (100 IU mL*^−^*^1^). After 11 days, cells were put in RPMI supplemented with 8 % human serum without any IL2. At day 12, cells were washed with RPMI, diluted at 2.10^6^ mL^-1^, and 200,000 cells plated in 96w round bottom plates. Then 100 uL of peptide pools were added (2 µM final in R8) and cells were incubated 1 h at 37°C. Protein Transport Inhibitor (1/1000, eBioscience 00-4980-93) was added and cells incubated for additional 4 h at 37°C. Cells were then washed with PBS and stained for 15 min at RT with fixable Near-IR Dead Cell Staining Kit (Thermofisher L10119, 1/1000 in PBS). After three washes, cells were stained with CD4 antibody (BD 562970) for 20 min at 4°C. After additional three washes, cells were incubated with Fix/perm kit (Biolegend 426803) for 20 min at 4°C in the dark, and stained with anti TNF*α* (BD 340512) and anti IFN*γ* (BD 554702) for 30 min at 4°C.

After final washing, cells were resuspended in FACS buffer (PBS 0.5 % FBS 2 mM EDTA) and analyzed on a Cytoflex S1 flow cytometer. Data were analyzed using the FlowJo v10.7.1 software. Positive and negative controls were obtained by incubating cells with PMA/ionomycin (Thermofisher, Cat# 00-4975-93) or without peptide, respectively. For peptide pools leading to an immune response, experiment was repeated with single peptides.

### Sorting of naives and effector/memory CD4^+^ T cells

Naive and effector/memory CD4^+^ T cells were isolated by Fluorescence-activated Cell Sorting (FACS) upon staining with anti-CD4 antibody (BD 562970), anti-CCR7 antibody (353227 BioLegend) and anti-CD45RA antibody (304108 BioLegend) for 30 min at 4°C. After three washes with FACS buffer (PBS 0.5 % FBS 2 mM EDTA) cells were incubated 10 min with DAPI (Sigma, Cat#10236276001) at 250 nM and washed again three times. Naïve (CCR7^+^ and CD45RA^+^) CD4 T cells and effector/memory (CD45RA^-^) CD4 T cells were collected separately.

### Peptide-MHC-II multimer validation and sorting of CD4^+^ T cells

CD4^+^ T cells were incubated with multimers (1/50 dilution) 45 min at 4°C in FACS buffer (PBS supplemented with 0.5 % FBS and 2 mM EDTA), isolated by FACS and either directly used for TCR sequencing or expanded with autologous irradiated CD4-depleted feeders in RPMI supplemented with 8 % human serum, phytohemagglutinin (Invivogen, 1 µg mL^-1^) and IL2 (150 IU mL^-1^).

### Bulk TCR sequencing

mRNA was extracted using the Dynabeads mRNA DIRECT purification kit according to the manufacturer instructions (ThermoFisher) and was then amplified using the MessageAmp II aRNA Amplification Kit (Ambion) with the following modifications: *in vitro* transcription was performed at 37°C for 16 h. First strand cDNA was synthesized using the Superscript III (Thermofisher) and a collection of TRAV/TRBV specific primers. Unique Molecular identifiers (UMI) of length 9 were added to each read. TCRs were then amplified by PCR (20 cycles with the Phusion from NEB) with a single primer pair binding to the constant region and the adapter linked to the TRAV/TRBV primers added during the reverse transcription. A second round of PCR (25 cycles with the Phusion from NEB) was performed to add the Illumina adapters containing the different indexes. The TCR products were purified with AMPure XP beads (Beckman Coulter), quantified and loaded on the MiniSeq instrument (Illumina) for deep sequencing of the TCRα/TCRβ chain.

### Analysis of TCR sequences

The fastq files were processed with MIGEC (Shugay et al., 2014), using default parameters to demultiplex them and identify the TCRα and TCRβ clonotypes. For each sample, the frequency of each TCR chain was computed based on UMI corrected counts. Only TCRs with more than one UMI count and representing more than 1% of the total UMI counts were considered. TCRs with the same amino acid sequences were merged in Table S3B.

The CDR3 sequences of the alpha and beta chains were used to search TCRα and TCRβ repertoires through the iReceptor web platform (Corrie et al., 2018), which contains, as of June 2022, 7,111 repertoires for a total of 5.1 billion sequences. Hits were defined as those having the same CDR3 sequence (Table S3B). We further restrict our analysis by considering only TCRs with the same CDR3 and the same V, J genes (100% sequence identity).

### Data and code availability

Mass spectrometry-based MHC-II peptidomics data generated for this study will be deposited in the ProteomeXchange Consortium via the PRIDE partner repository (Perez-Riverol et al., 2022).

The models of the crystal structures resolved in this work will be deposited in the worldwide Protein Data Bank (Berman et al., 2003).

TCR sequencing data will be deposited in NCBI’s Gene Expression Omnibus (Edgar et al., 2002).

MixMHC2pred is freely available for academic usage (https://github.com/GfellerLab/MixMHC2pred) and it is also available through a webserver (http://mixmhc2pred.gfellerlab.org).

## Notes

### Competing Interest Statement

The authors have declared no competing interest.

## References

1. Abelin, J.G., Harjanto, D., Malloy, M., Suri, P., Colson, T., Goulding, S.P., Creech, A.L., Serrano, L.R., Nasir, G., Nasrullah, Y., et al. (2019). Defining HLA-II Ligand Processing and Binding Rules with Mass Spectrometry Enhances Cancer Epitope Prediction. Immunity 51, 766–779.e17.

2. Afrache, H., Tregaskes, C.A., and Kaufman, J. (2020). A potential nomenclature for the Immuno Polymorphism Database (IPD) of chicken MHC genes: progress and problems. Immunogenetics 72, 9–24.

3. Alspach, E., Lussier, D.M., Miceli, A.P., Kizhvatov, I., DuPage, M., Luoma, A.M., Meng, W., Lichti, C.F., Esaulova, E., Vomund, A.N., et al. (2019). MHC-II neoantigens shape tumour immunity and response to immunotherapy. Nature 574, 696–701.

4. Balen, P. van, Kester, M.G.D., Klerk, W. de, Crivello, P., Arrieta-Bolaños, E., Ru, A.H. de, Jedema, I., Mohammed, Y., Heemskerk, M.H.M., Fleischhauer, K., et al. (2020). Immunopeptidome Analysis of HLA-DPB1 Allelic Variants Reveals New Functional Hierarchies. J. Immunol. 204, 3273–3282.

5. Barra, C., Alvarez, B., Paul, S., Sette, A., Peters, B., Andreatta, M., Buus, S., and Nielsen, M. (2018). Footprints of antigen processing boost MHC class II natural ligand predictions. Genome Med. 10, 84.

6. Bergseng, E., Dørum, S., Arntzen, M.Ø., Nielsen, M., Nygård, S., Buus, S., Souza, G.A. de, and Sollid, L.M. (2015). Different binding motifs of the celiac disease-associated HLA molecules DQ2.5, DQ2.2, and DQ7.5 revealed by relative quantitative proteomics of endogenous peptide repertoires. Immunogenetics 67, 73–84.

7. Berman, H., Henrick, K., and Nakamura, H. (2003). Announcing the worldwide Protein Data Bank. Nat. Struct. Mol. Biol. 10, 980–980.

8. Berman, H.M., Westbrook, J., Feng, Z., Gilliland, G., Bhat, T.N., Weissig, H., Shindyalov, I.N., and Bourne, P.E. (2000). The Protein Data Bank. Nucleic Acids Res. 28, 235–242.

9. Bird, P.I., Trapani, J.A., and Villadangos, J.A. (2009). Endolysosomal proteases and their inhibitors in immunity. Nat. Rev. Immunol. 9, 871–882.

10. Bodenhofer, U., Bonatesta, E., Horejš-Kainrath, C., and Hochreiter, S. (2015). msa: an R package for multiple sequence alignment. Bioinformatics 31, 3997–3999.

11. Borst, J., Ahrends, T., B*ą*ba*ł*a, N., Melief, C.J.M., and Kastenmüller, W. (2018). CD4+ T cell help in cancer immunology and immunotherapy. Nat. Rev. Immunol. 18, 635–647.

12. Cassotta, A., Paparoditis, P., Geiger, R., Mettu, R.R., Landry, S.J., Donati, A., Benevento, M., Foglierini, M., Lewis, D.J.M., Lanzavecchia, A., et al. (2020). Deciphering and predicting CD4+ T cell immunodominance of influenza virus hemagglutinin. J. Exp. Med. 217.

13. Chen, B., Khodadoust, M.S., Olsson, N., Wagar, L.E., Fast, E., Liu, C.L., Muftuoglu, Y., Sworder, B.J., Diehn, M., Levy, R., et al. (2019). Predicting HLA class II antigen presentation through integrated deep learning. Nat. Biotechnol. 1–12.

14. Ciudad, M.T., Sorvillo, N., Alphen, F.P. van, Catalán, D., Meijer, A.B., Voorberg, J., and Jaraquemada, D. (2017). Analysis of the HLA-DR peptidome from human dendritic cells reveals high affinity repertoires and nonconventional pathways of peptide generation. J. Leukoc. Biol. 101, 15–27.

15. Clement, C.C., Becerra, A., Yin, L., Zolla, V., Huang, L., Merlin, S., Follenzi, A., Shaffer, S.A., Stern, L.J., and Santambrogio, L. (2016). The Dendritic Cell Major Histocompatibility Complex II (MHC II) Peptidome Derives from a Variety of Processing Pathways and Includes Peptides with a Broad Spectrum of HLA-DM Sensitivity. J. Biol. Chem. 291, 5576–5595.

16. Collado, J.A., Alvarez, I., Ciudad, M.T., Espinosa, G., Canals, F., Pujol-Borrell, R., Carrascal, M., Abian, J., and Jaraquemada, D. (2013). Composition of the HLA-DR-associated human thymus peptidome. Eur. J. Immunol. 43, 2273–2282.

17. Corrie, B.D., Marthandan, N., Zimonja, B., Jaglale, J., Zhou, Y., Barr, E., Knoetze, N., Breden, F.M.W., Christley, S., Scott, J.K., et al. (2018). iReceptor: A platform for querying and analyzing antibody/B-cell and T-cell receptor repertoire data across federated repositories. Immunol. Rev. 284, 24–41.

18. Cox, J., and Mann, M. (2008). MaxQuant enables high peptide identification rates, individualized p.p.b.-range mass accuracies and proteome-wide protein quantification. Nat. Biotechnol. 26, 1367–1372.

19. Deutsch, E.W., Bandeira, N., Sharma, V., Perez-Riverol, Y., Carver, J.J., Kundu, D.J., García-Seisdedos, D., Jarnuczak, A.F., Hewapathirana, S., Pullman, B.S., et al. (2020). The ProteomeXchange consortium in 2020: enabling ‘big data’ approaches in proteomics. Nucleic Acids Res. 48, D1145–D1152.

20. Dheilly, E., Battistello, E., Katanayeva, N., Sungalee, S., Michaux, J., Duns, G., Wehrle, S., Sordet-Dessimoz, J., Mina, M., Racle, J., et al. (2020). Cathepsin S Regulates Antigen Processing and T Cell Activity in Non-Hodgkin Lymphoma. Cancer Cell 37, 674–689.e12.

21. Draheim, M., Wlodarczyk, M.F., Crozat, K., Saliou, J.-M., Alayi, T.D., Tomavo, S., Hassan, A., Salvioni, A., Demarta-Gatsi, C., Sidney, J., et al. (2017). Profiling MHC II immunopeptidome of blood-stage malaria reveals that cDC1 control the functionality of parasite-specific CD4 T cells. EMBO Mol. Med. 9, 1605–1621.

22. Edgar, R., Domrachev, M., and Lash, A.E. (2002). Gene Expression Omnibus: NCBI gene expression and hybridization array data repository. Nucleic Acids Res. 30, 207–210.

23. Emsley, P., and Cowtan, K. (2004). Coot: model-building tools for molecular graphics. Acta Crystallogr. D Biol. Crystallogr. 60, 2126–2132.

24. Falk, K., Rötzschke, O., Stevanovíc, S., Jung, G., and Rammensee, H.-G. (1994). Pool sequencing of natural HLA-DR, DQ, and DP ligands reveals detailed peptide motifs, constraints of processing, and general rules. Immunogenetics 39, 230–242.

25. Fisch, A., Reynisson, B., Benedictus, L., Nicastri, A., Vasoya, D., Morrison, I., Buus, S., Ferreira, B.R., Santos, I.K.F. de M., Ternette, N., et al. (2021). Integral Use of Immunopeptidomics and Immunoinformatics for the Characterization of Antigen Presentation and Rational Identification of BoLA-DR–Presented Peptides and Epitopes. J. Immunol.

26. Forlani, G., Michaux, J., Pak, H., Huber, F., Joseph, E.L.M., Ramia, E., Stevenson, B.J., Linnebacher, M., Accolla, R., and Bassani-Sternberg, M. (2020). CIITA-transduced glioblastoma cells uncover a rich repertoire of clinically relevant tumor-associated HLA-II antigens. Mol. Cell. Proteomics.

27. Garde, C., Ramarathinam, S.H., Jappe, E.C., Nielsen, M., Kringelum, J.V., Trolle, T., and Purcell, A.W. (2019). Improved peptide-MHC class II interaction prediction through integration of eluted ligand and peptide affinity data. Immunogenetics 71, 445–454.

28. Goncalves, G., Mullan, K.A., Duscharla, D., Ayala, R., Croft, N.P., Faridi, P., and Purcell, A.W. (2021). IFN*γ* Modulates the Immunopeptidome of Triple Negative Breast Cancer Cells by Enhancing and Diversifying Antigen Processing and Presentation. Front. Immunol. 12.

29. Graciotti, M., Marino, F., Pak, H., Baumgaertner, P., Thierry, A.-C., Chiffelle, J., Perez, M.A.S., Zoete, V., Harari, A., Bassani-Sternberg, M., et al. (2020). Deciphering the Mechanisms of Improved Immunogenicity of Hypochlorous Acid-Treated Antigens in Anti-Cancer Dendritic Cell-Based Vaccines. Vaccines 8, 271.

30. Greenbaum, J., Sidney, J., Chung, J., Brander, C., Peters, B., and Sette, A. (2011). Functional classification of class II human leukocyte antigen (HLA) molecules reveals seven different supertypes and a surprising degree of repertoire sharing across supertypes. Immunogenetics 63, 325–335.

31. Günther, S., Schlundt, A., Sticht, J., Roske, Y., Heinemann, U., Wiesmüller, K.-H., Jung, G., Falk, K., Rötzschke, O., and Freund, C. (2010). Bidirectional binding of invariant chain peptides to an MHC class II molecule. Proc. Natl. Acad. Sci. 107, 22219–22224.

32. Halabi, S., Ghosh, M., Stevanović, S., Rammensee, H.-G., Bertzbach, L.D., Kaufer, B.B., Moncrieffe, M.C., Kaspers, B., Härtle, S., and Kaufman, J. (2021). The dominantly expressed class II molecule from a resistant MHC haplotype presents only a few Marek’s disease virus peptides by using an unprecedented binding motif. PLOS Biol. 19, e3001057.

33. Henikoff, S., and Henikoff, J.G. (1992). Amino acid substitution matrices from protein blocks. Proc. Natl. Acad. Sci. 89, 10915–10919.

34. Holland, C.J., Cole, D.K., and Godkin, A. (2013). Re-Directing CD4+ T Cell Responses with the Flanking Residues of MHC Class II-Bound Peptides: The Core is Not Enough. Front. Immunol. 4.

35. Hu, Z., Leet, D.E., Allesøe, R.L., Oliveira, G., Li, S., Luoma, A.M., Liu, J., Forman, J., Huang, T., Iorgulescu, J.B., et al. (2021). Personal neoantigen vaccines induce persistent memory T cell responses and epitope spreading in patients with melanoma. Nat. Med. 1–11.

36. Kabsch, W. (2010). XDS. Acta Crystallogr. D Biol. Crystallogr. 66, 125–132.

37. Kalaora, S., Nagler, A., Nejman, D., Alon, M., Barbolin, C., Barnea, E., Ketelaars, S.L.C., Cheng, K., Vervier, K., Shental, N., et al. (2021). Identification of bacteria-derived HLA-bound peptides in melanoma. Nature 1–6.

38. Khodadoust, M.S., Olsson, N., Wagar, L.E., Haabeth, O.A.W., Chen, B., Swaminathan, K., Rawson, K., Liu, C.L., Steiner, D., Lund, P., et al. (2017). Antigen presentation profiling reveals recognition of lymphoma immunoglobulin neoantigens. Nature 543, 723– 727.

39. Laghmouchi, A., Kester, M.G.D., Hoogstraten, C., Hageman, L., de Klerk, W., Huisman, W., Koster, E.A.S., de Ru, A.H., van Balen, P., Klobuch, S., et al. (2022). Promiscuity of Peptides Presented in HLA-DP Molecules from Different Immunogenicity Groups Is Associated With T-Cell Cross-Reactivity. Front. Immunol. 13.

40. Liebschner, D., Afonine, P.V., Baker, M.L., Bunkóczi, G., Chen, V.B., Croll, T.I., Hintze, B., Hung, L.-W., Jain, S., McCoy, A.J., et al. (2019). Macromolecular structure determination using X-rays, neutrons and electrons: recent developments in Phenix. Acta Crystallogr. Sect. Struct. Biol. 75, 861–877.

41. Maccari, G., Robinson, J., Ballingall, K., Guethlein, L.A., Grimholt, U., Kaufman, J., Ho, C.-S., de Groot, N.G., Flicek, P., Bontrop, R.E., et al. (2017). IPD-MHC 2.0: an improved inter-species database for the study of the major histocompatibility complex. Nucleic Acids Res. 45, D860–D864.

42. Marcu, A., Bichmann, L., Kuchenbecker, L., Kowalewski, D.J., Freudenmann, L.K., Backert, L., Mühlenbruch, L., Szolek, A., Lübke, M., Wagner, P., et al. (2021). HLA Ligand Atlas: a benign reference of HLA-presented peptides to improve T-cell-based cancer immunotherapy. J. Immunother. Cancer 9, e002071.

43. Marino, F., Semilietof, A., Michaux, J., Pak, H.-S., Coukos, G., Müller, M., and Bassani-Sternberg, M. (2020). Biogenesis of HLA Ligand Presentation in Immune Cells Upon Activation Reveals Changes in Peptide Length Preference. Front. Immunol. 11.

44. Meurer, T., Crivello, P., Metzing, M., Kester, M., Megger, D.A., Chen, W., van Veelen, P.A., van Balen, P., Westendorf, A.M., Homa, G., et al. (2021). Permissive HLA-DPB1 mismatches in HCT depend on immunopeptidome divergence and editing by HLA-DM. Blood 137, 923–928.

45. Neefjes, J., Jongsma, M.L.M., Paul, P., and Bakke, O. (2011). Towards a systems understanding of MHC class I and MHC class II antigen presentation. Nat. Rev. Immunol. 11, 823–836.

46. Nelde, A., Kowalewski, D.J., Backert, L., Schuster, H., Werner, J.-O., Klein, R., Kohlbacher, O., Kanz, L., Salih, H.R., Rammensee, H.-G., et al. (2018). HLA ligandome analysis of primary chronic lymphocytic leukemia (CLL) cells under lenalidomide treatment confirms the suitability of lenalidomide for combination with T-cell-based immunotherapy. OncoImmunology 7, e1316438.

47. Newey, A., Griffiths, B., Michaux, J., Pak, H.S., Stevenson, B.J., Woolston, A., Semiannikova, M., Spain, G., Barber, L.J., Matthews, N., et al. (2019). Immunopeptidomics of colorectal cancer organoids reveals a sparse HLA class I neoantigen landscape and no increase in neoantigens with interferon or MEK-inhibitor treatment. J. Immunother. Cancer 7, 309.

48. Nielsen, M., Lundegaard, C., Worning, P., Hvid, C.S., Lamberth, K., Buus, S., Brunak, S., and Lund, O. (2004). Improved prediction of MHC class I and class II epitopes using a novel Gibbs sampling approach. Bioinformatics 20, 1388–1397.

49. Ooi, J.D., Petersen, J., Tan, Y.H., Huynh, M., Willett, Z.J., Ramarathinam, S.H., Eggenhuizen, P.J., Loh, K.L., Watson, K.A., Gan, P.Y., et al. (2017). Dominant protection from HLA-linked autoimmunity by antigen-specific regulatory T cells. Nature 545, 243–247.

50. Ott, P.A., Hu, Z., Keskin, D.B., Shukla, S.A., Sun, J., Bozym, D.J., Zhang, W., Luoma, A., Giobbie-Hurder, A., Peter, L., et al. (2017). An immunogenic personal neoantigen vaccine for patients with melanoma. Nature 547, 217–221.

51. Paul, S., Karosiene, E., Dhanda, S.K., Jurtz, V., Edwards, L., Nielsen, M., Sette, A., and Peters, B. (2018). Determination of a Predictive Cleavage Motif for Eluted Major Histocompatibility Complex Class II Ligands. Front. Immunol. 9.

52. Perez-Riverol, Y., Bai, J., Bandla, C., García-Seisdedos, D., Hewapathirana, S., Kamatchinathan, S., Kundu, D.J., Prakash, A., Frericks-Zipper, A., Eisenacher, M., et al. (2022). The PRIDE database resources in 2022: a hub for mass spectrometry-based proteomics evidences. Nucleic Acids Res. 50, D543–D552.

53. Racle, J., Michaux, J., Rockinger, G.A., Arnaud, M., Bobisse, S., Chong, C., Guillaume, P., Coukos, G., Harari, A., Jandus, C., et al. (2019). Robust prediction of HLA class II epitopes by deep motif deconvolution of immunopeptidomes. Nat. Biotechnol. 37, 1283–1286.

54. Ramarathinam, S.H., Ho, B.K., Dudek, N.L., and Purcell, A.W. (2021). HLA class II immunopeptidomics reveals that co-inherited HLA-allotypes within an extended haplotype can improve proteome coverage for immunosurveillance. PROTEOMICS 21, 2000160.

55. Reynisson, B., Barra, C., Kaabinejadian, S., Hildebrand, W.H., Peters, B., and Nielsen, M. (2020). Improved Prediction of MHC II Antigen Presentation through Integration and Motif Deconvolution of Mass Spectrometry MHC Eluted Ligand Data. J. Proteome Res. 19, 2304–2315.

56. Ritz, D., Sani, E., Debiec, H., Ronco, P., Neri, D., and Fugmann, T. (2018). Membranal and Blood-Soluble HLA Class II Peptidome Analyses Using Data-Dependent and Independent Acquisition. PROTEOMICS 18, 1700246.

57. Robinson, J., Barker, D.J., Georgiou, X., Cooper, M.A., Flicek, P., and Marsh, S.G.E. (2020). IPD-IMGT/HLA Database. Nucleic Acids Res. 48, D948–D955.

58. Rose, P.W., Prlić, A., Altunkaya, A., Bi, C., Bradley, A.R., Christie, C.H., Costanzo, L.D., Duarte, J.M., Dutta, S., Feng, Z., et al. (2017). The RCSB protein data bank: integrative view of protein, gene and 3D structural information. Nucleic Acids Res. 45, D271–D281.

59. Sahin, U., Derhovanessian, E., Miller, M., Kloke, B.-P., Simon, P., Löwer, M., Bukur, V., Tadmor, A.D., Luxemburger, U., Schrörs, B., et al. (2017). Personalized RNA mutanome vaccines mobilize poly-specific therapeutic immunity against cancer. Nature 547, 222–226.

60. Schlundt, A., Günther, S., Sticht, J., Wieczorek, M., Roske, Y., Heinemann, U., and Freund, C. (2012). Peptide Linkage to the *α*-Subunit of MHCII Creates a Stably Inverted Antigen Presentation Complex. J. Mol. Biol. 423, 294–302.

61. Schymkowitz, J., Borg, J., Stricher, F., Nys, R., Rousseau, F., and Serrano, L. (2005). The FoldX web server: an online force field. Nucleic Acids Res. 33, W382–W388.

62. Sercarz, E.E., and Maverakis, E. (2003). MHC-guided processing: binding of large antigen fragments. Nat. Rev. Immunol. 3, 621–629.

63. Shao, X.M., Bhattacharya, R., Huang, J., Sivakumar, I.K.A., Tokheim, C., Zheng, L., Hirsch, D., Kaminow, B., Omdahl, A., Bonsack, M., et al. (2020). High-Throughput Prediction of MHC Class I and II Neoantigens with MHCnuggets. Cancer Immunol. Res. 8, 396–408.

64. Shugay, M., Britanova, O.V., Merzlyak, E.M., Turchaninova, M.A., Mamedov, I.Z., Tuganbaev, T.R., Bolotin, D.A., Staroverov, D.B., Putintseva, E.V., Plevova, K., et al. (2014). Towards error-free profiling of immune repertoires. Nat. Methods 11, 653–655.

65. Sievers, F., Wilm, A., Dineen, D., Gibson, T.J., Karplus, K., Li, W., Lopez, R., McWilliam, H., Remmert, M., Söding, J., et al. (2011). Fast, scalable generation of high-quality protein multiple sequence alignments using Clustal Omega. Mol. Syst. Biol. 7, 539.

66. Sofron, A., Ritz, D., Neri, D., and Fugmann, T. (2016). High-resolution analysis of the murine MHC class II immunopeptidome. Eur. J. Immunol. 46, 319–328.

67. The UniProt Consortium (2021). UniProt: the universal protein knowledgebase in 2021. Nucleic Acids Res. 49, D480–D489.

68. Ting, Y.T., Petersen, J., Ramarathinam, S.H., Scally, S.W., Loh, K.L., Thomas, R., Suri, A., Baker, D.G., Purcell, A.W., Reid, H.H., et al. (2018). The interplay between citrullination and HLA-DRB1 polymorphism in shaping peptide binding hierarchies in rheumatoid arthritis. J. Biol. Chem. 293, 3236–3251.

69. Tran, E., Turcotte, S., Gros, A., Robbins, P.F., Lu, Y.-C., Dudley, M.E., Wunderlich, J.R., Somerville, R.P., Hogan, K., Hinrichs, C.S., et al. (2014). Cancer Immunotherapy Based on Mutation-Specific CD4+ T Cells in a Patient with Epithelial Cancer. Science 344, 641–645.

70. Unanue, E.R., Turk, V., and Neefjes, J. (2016). Variations in MHC Class II Antigen Processing and Presentation in Health and Disease. Annu. Rev. Immunol. 34, 265–297.

71. Vita, R., Mahajan, S., Overton, J.A., Dhanda, S.K., Martini, S., Cantrell, J.R., Wheeler, D.K., Sette, A., and Peters, B. (2019). The Immune Epitope Database (IEDB): 2018 update. Nucleic Acids Res. 47, D339–D343.

72. Wagih, O. (2017). ggseqlogo: a versatile R package for drawing sequence logos. Bioinformatics 33, 3645–3647.

73. Wan, X., Vomund, A.N., Peterson, O.J., Chervonsky, A.V., Lichti, C.F., and Unanue, E.R. (2020). The MHC-II peptidome of pancreatic islets identifies key features of autoimmune peptides. Nat. Immunol. 1–9.

74. Wang, Q., Drouin, E.E., Yao, C., Zhang, J., Huang, Y., Leon, D.R., Steere, A.C., and Costello, C.E. (2017). Immunogenic HLA-DR-Presented Self-Peptides Identified Directly from Clinical Samples of Synovial Tissue, Synovial Fluid, or Peripheral Blood in Patients with Rheumatoid Arthritis or Lyme Arthritis. J. Proteome Res. 16, 122–136.

75. Webb, B., and Sali, A. (2016). Comparative Protein Structure Modeling Using MODELLER. Curr. Protoc. Bioinforma. 54, 5.6.1–5.6.37.

76. Zacharakis, N., Chinnasamy, H., Black, M., Xu, H., Lu, Y.-C., Zheng, Z., Pasetto, A., Langhan, M., Shelton, T., Prickett, T., et al. (2018). Immune recognition of somatic mutations leading to complete durable regression in metastatic breast cancer. Nat. Med. 24, 724–730.

