## Supplementary figures and tables for "Machine learning predictions of MHC-II specificities reveal alternative binding mode of class II epitopes"

### Figure S1. A

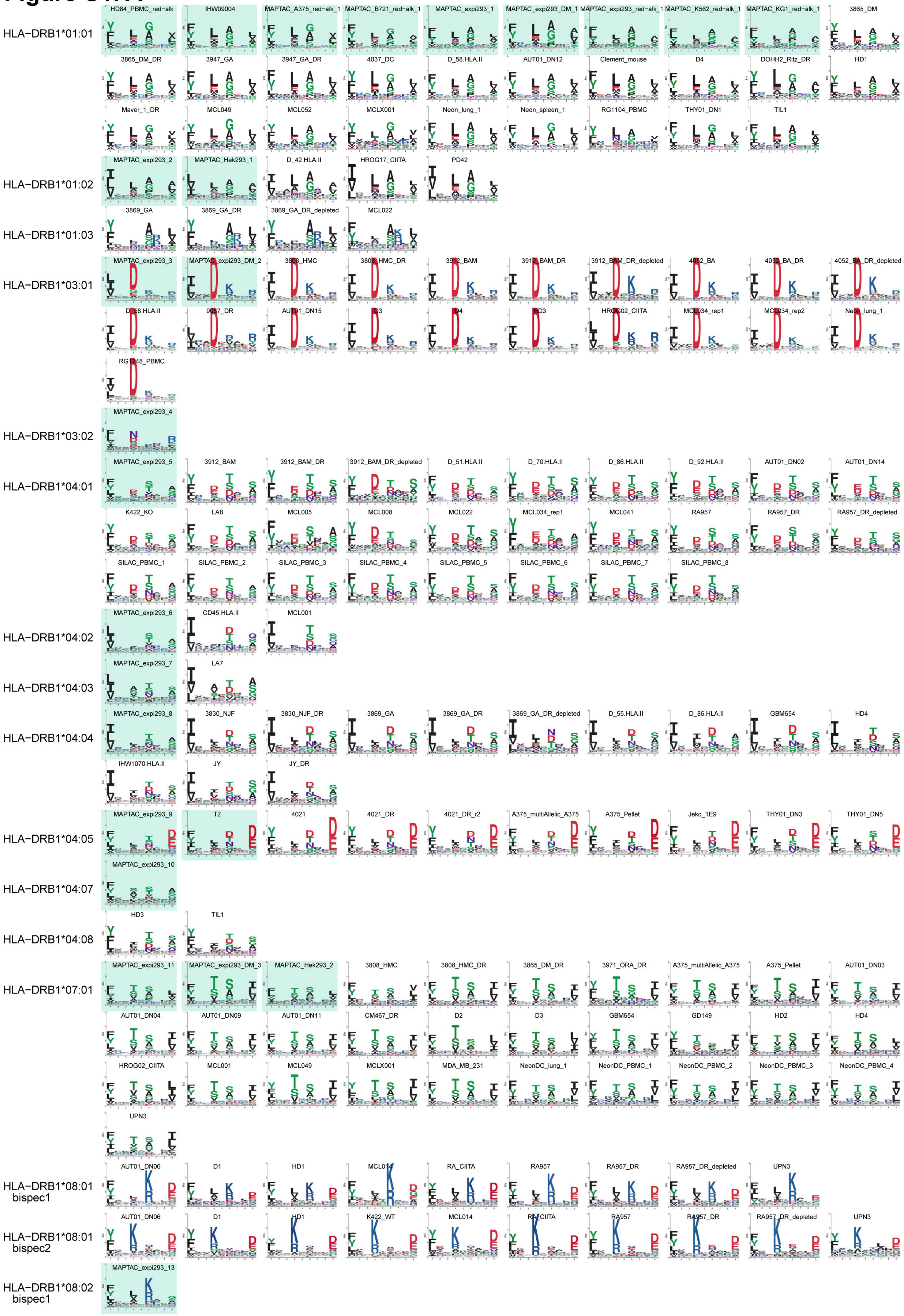

HLA-DRB1*01:01
HD84_PBMC_red-alk
IHW09004
MAPTAC_A375_red-alk_1
MAPTAC_B721_red-alk_1
MAPTAC_expi293_1
MAPTAC_expi293_DM_1
MAPTAC_expi293_red-alk_1
MAPTAC_K562_red-alk_1
MAPTAC_KG1_red-alk_1
3865_DM
3865_DM_DR
3947_GA
3947_GA_DR
4037_DC
D_58.HLA.II
AUT01_DN12
Clement_mouse
D4
DOHH2_Ritz_DR
HD1
Maver_1_DR
MCL049
MCL052
MCLX001
Neon_lung_1
Neon_spleen_1
RG1104_PBMC
THY01_DN1
TIL1
HLA-DRB1*01:02
MAPTAC_expi293_2
MAPTAC_Hek293_1
D_42.HLA.II
HROG17_CIITA
PD42
HLA-DRB1*01:03
3869_GA
3869_GA_DR
3869_GA_DR_depleted
MCL022
HLA-DRB1*03:01
MAPTAC_expi293_3
MAPTAC_expi293_DM_2
3808_HMC
3808_HMC_DR
3912_BAM
3912_BAM_DR
3912_BAM_DR_depleted
4052_BA
4052_BA_DR
4052_BA_DR_depleted
D_58.HLA.II
9087_DR
AUT01_DN15
D3
D4
HD3
HROG02_CIITA
MCL034_rep1
MCL034_rep2
Neon_lung_1
RG1248_PBMC
HLA-DRB1*03:02
MAPTAC_expi293_4
HLA-DRB1*04:01
MAPTAC_expi293_5
3912_BAM
3912_BAM_DR
3912_BAM_DR_depleted
D_51.HLA.II
D_70.HLA.II
D_86.HLA.II
D_92.HLA.II
AUT01_DN02
AUT01_DN14
K422_KO
LA8
MCL005
MCL008
MCL022
MCL034_rep1
MCL041
RA957
RA957_DR
RA957_DR_depleted
SILAC_PBMC_1
SILAC_PBMC_2
SILAC_PBMC_3
SILAC_PBMC_4
SILAC_PBMC_5
SILAC_PBMC_6
SILAC_PBMC_7
SILAC_PBMC_8
HLA-DRB1*04:02
MAPTAC_expi293_6
CD45.HLA.II
MCL001
HLA-DRB1*04:03
MAPTAC_expi293_7
LA7
HLA-DRB1*04:04
MAPTAC_expi293_8
3830_NJF
3830_NJF_DR
3869_GA
3869_GA_DR
3869_GA_DR_depleted
D_55.HLA.II
D_86.HLA.II
GBM654
HD4
IHW1070.HLA.II
JY
JY_DR
HLA-DRB1*04:05
MAPTAC_expi293_9
T2
4021
4021_DR
4021_DR_r2
A375_multiAllelic_A375
A375_Pellet
Jeko_1E9
THY01_DN3
THY01_DN5
HLA-DRB1*04:07
MAPTAC_expi293_10
HLA-DRB1*04:08
HD3
TIL1
HLA-DRB1*07:01
MAPTAC_expi293_11
MAPTAC_expi293_DM_3
MAPTAC_Hek293_2
3808_HMC
3808_HMC_DR
3865_DM_DR
3971_ORA_DR
A375_multiAllelic_A375
A375_Pellet
AUT01_DN03
AUT01_DN04
AUT01_DN09
AUT01_DN11
CM467_DR
D2
D3
GBM654
GD149
HD2
HD4
HROG02_CIITA
MCL001
MCL049
MCLX001
MDA_MB_231
NeonDC_lung_1
NeonDC_PBMC_1
NeonDC_PBMC_2
NeonDC_PBMC_3
NeonDC_PBMC_4
UPN3
HLA-DRB1*08:01 bispec1
AUT01_DN06
D1
HD1
MCL014
RA_CIITA
RA957
RA957_DR
RA957_DR_depleted
UPN3
HLA-DRB1*08:01 bispec2
AUT01_DN06
D1
HD1
K422_WT
MCL014
RA_CIITA
RA957
RA957_DR
RA957_DR_depleted
UPN3
HLA-DRB1*08:02 bispec1
MAPTAC_expi293_13

### Figure S1. A (continued)

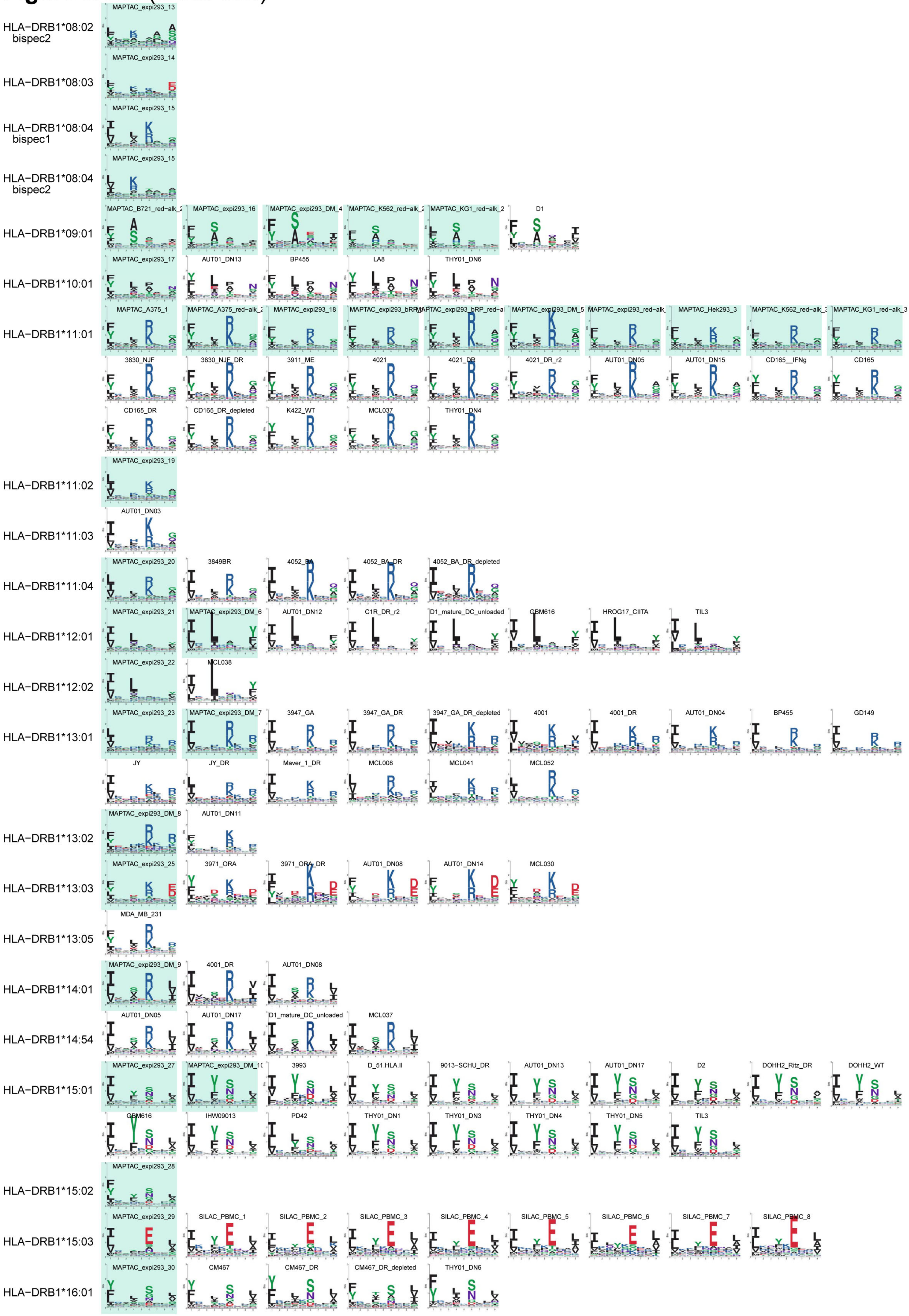

### Figure S1. A (continued)

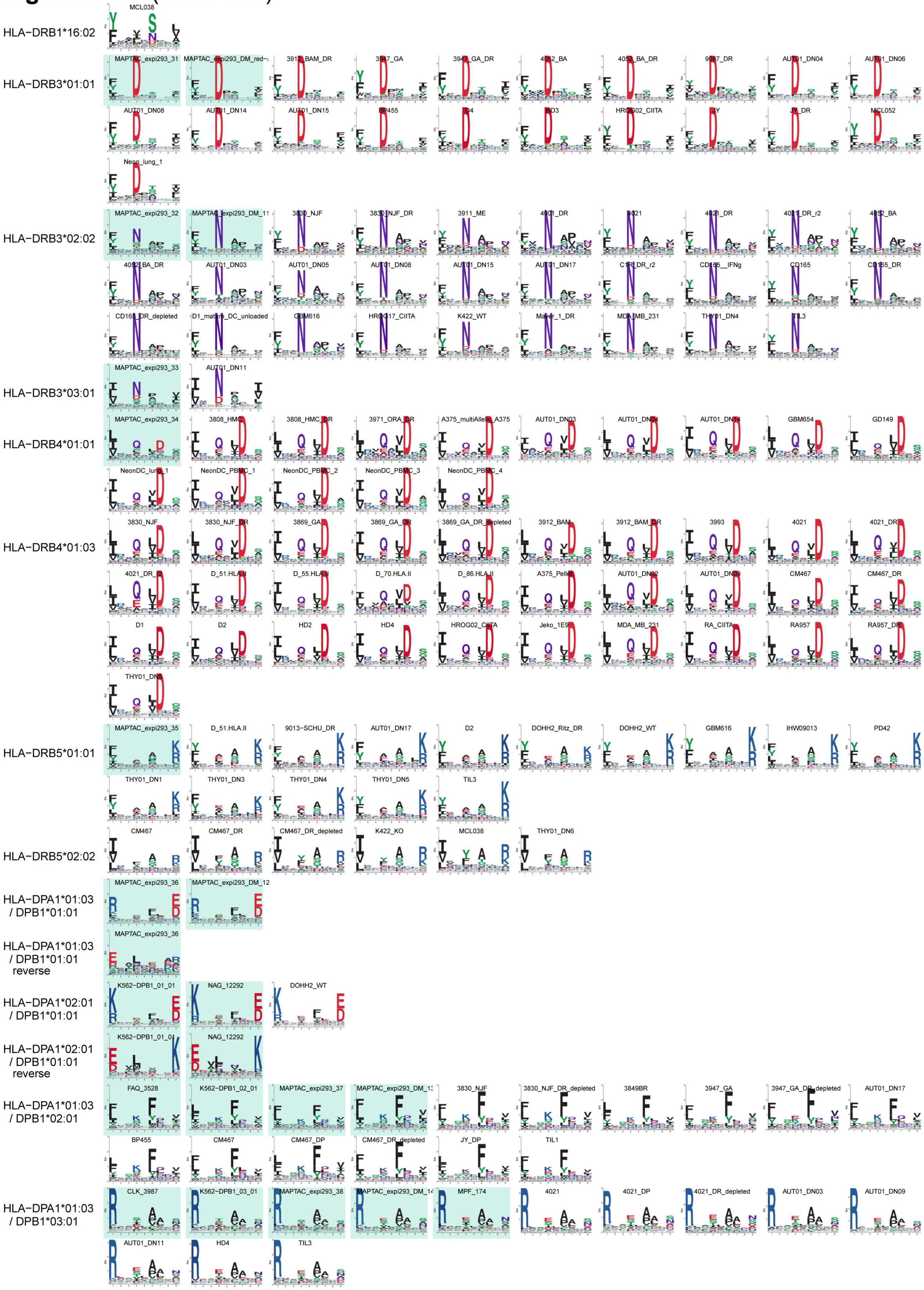

### Figure S1. A (continued)

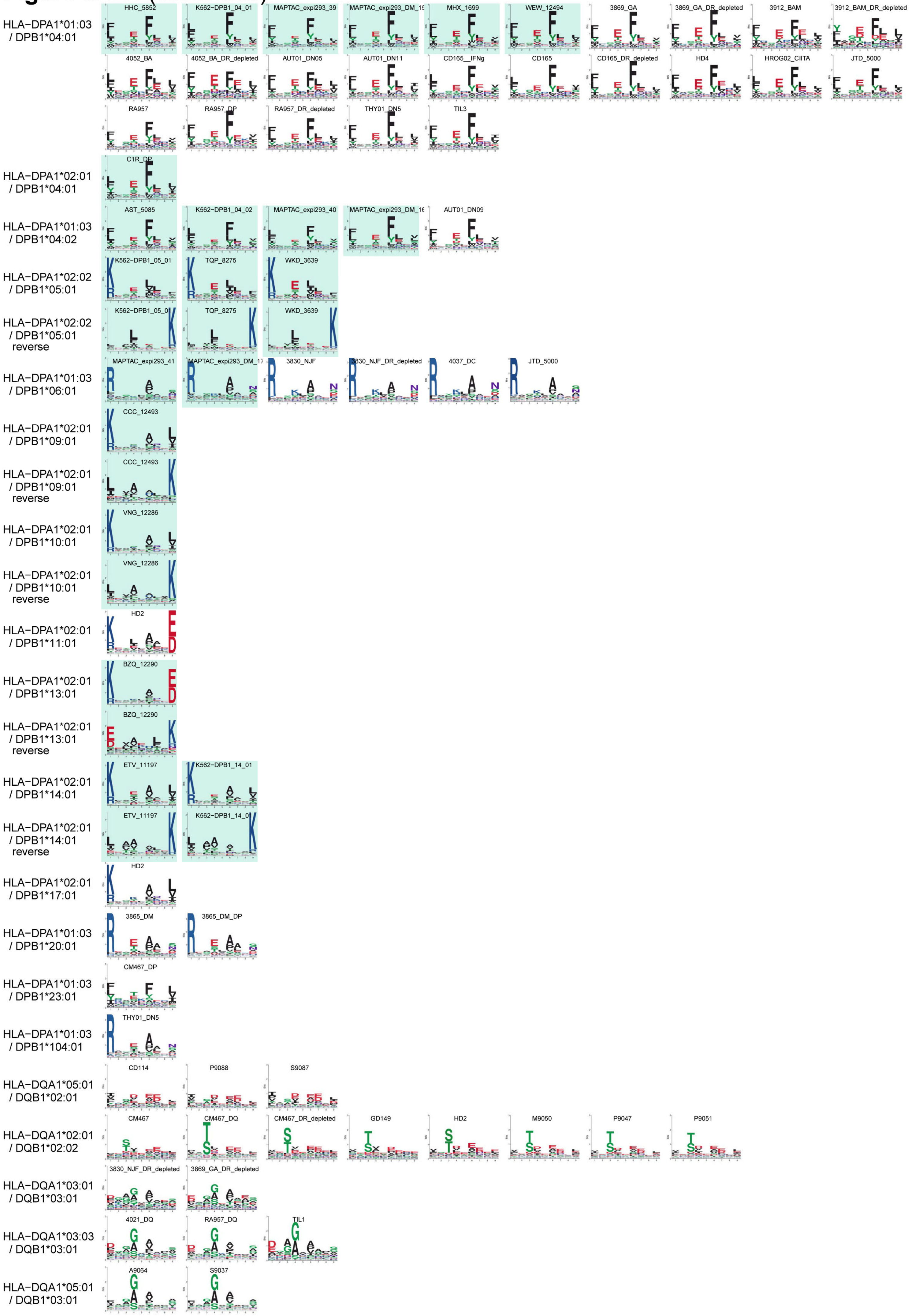

**Figure S1. A** (continued)

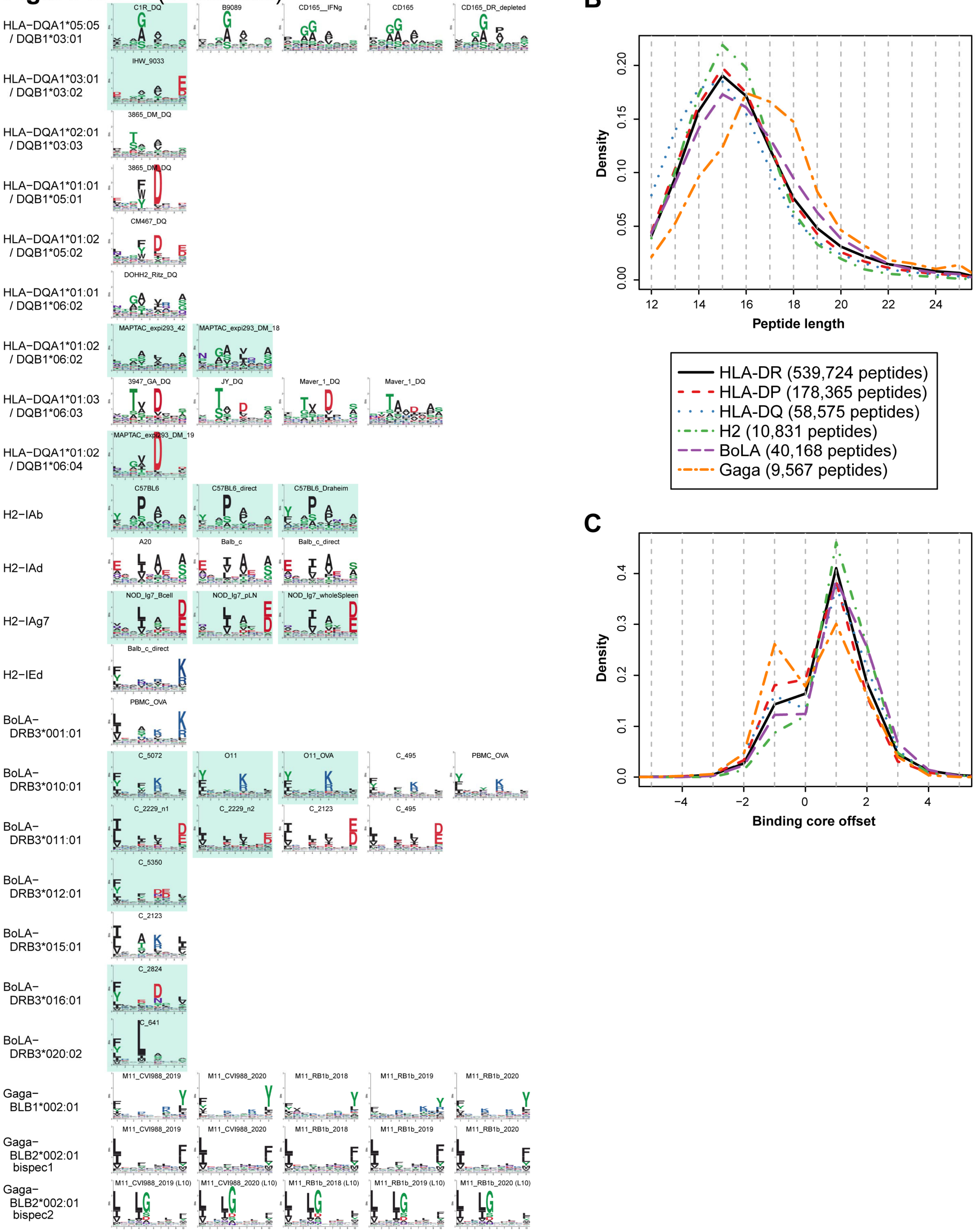

**Figure S1. Curation of MHC-II peptidomics data reveals binding specificities for 88 MHC-II alleles.**

(A) MHC-II binding motifs obtained with MoDec for each allele across all MHC-II peptidomics samples where the motif could be accurately resolved. The name of the sample is indicated above each motif. Motifs with a light-green background are coming from monoallelic samples.

(B) Peptide length distributions per allele class and species. The number of peptides indicated in the legend includes duplicate peptides (same peptide found in multiple samples).

(C) Peptide binding core offset distribution per allele class and species.

The more evolutionary distant chicken alleles follow slightly different distributions than other MHC-II alleles, in line with the longer binding core that they can accommodate.

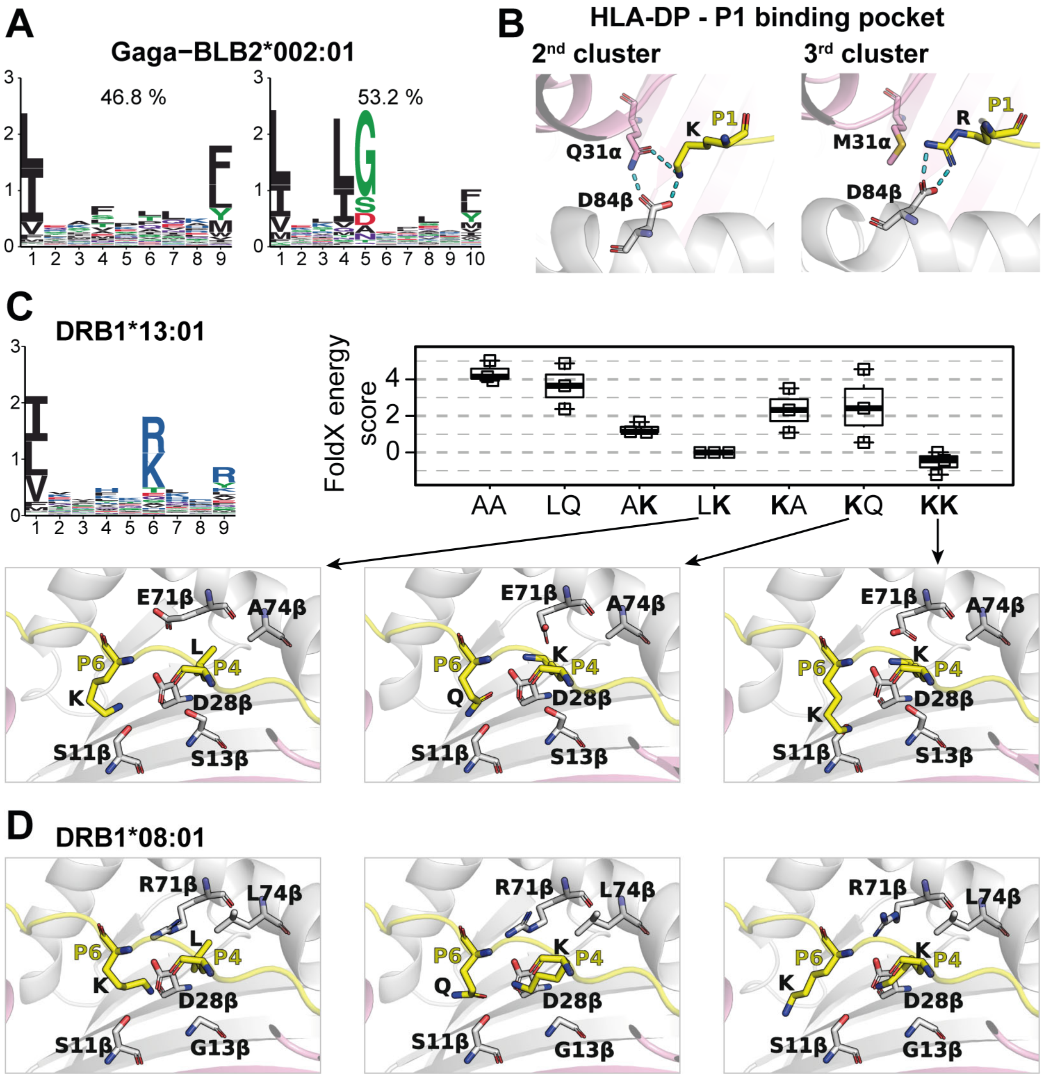


**Figure S2. MHC-II binding specificities reflect biochemical properties of the MHC-II binding pockets.**

(A) Motifs observed for Gaga-BLB2*002:01 allele, where some ligands are described by a 9-mer binding motif and other by a 10-mer binding motif. Percentage above the motifs indicate the fraction of peptides assigned to each sub-specificity (based on 12,289 peptides in total, including duplicates).

(B) Polar and charged interactions (showed in cyan) formed between residues in the MHC-II binding site and the residue K/R found at P1 in HLA-DP ligands from the $2^{nd}$ and $3^{rd}$ clusters of Figure 2A.

(C) Binding motif (top-left), FoldX energy score (top-right) and structural modeling (bottom) of peptides in complex with HLA-DRB1*13:01. A single binding specificity is observed for this allele in MHC-II peptidomics data. The boxplot shows the calculated change in the FoldX energy score for various peptides with 0, 1 or 2 positively charged AAs at P4 and P6 (simulations based on three different peptides for each case, differences in FoldX energy score are relative to the "LK" case). The bottom part shows a model of HLA-DRB1*13:01 with multiple residues from the alleles that can interact with the peptide P4 and P6 binding core positions.

(D) Similar structural model for HLA-DRB1*08:01 shows that the main differences with HLA-DRB1*13:01 around the P4-P6 peptide binding anchors lie in the residues 13β (G in HLA-DRB1*08:01 and S in HLA-DRB1*13:01) and 71β (R in HLA-DRB1*08:01 and E in HLA-DRB1*13:01) that make that a K at P4 from the peptide will either interact with D28β (HLA-DRB1*08:01) or E71β (HLA-DRB1*13:01).

#### Figure S3

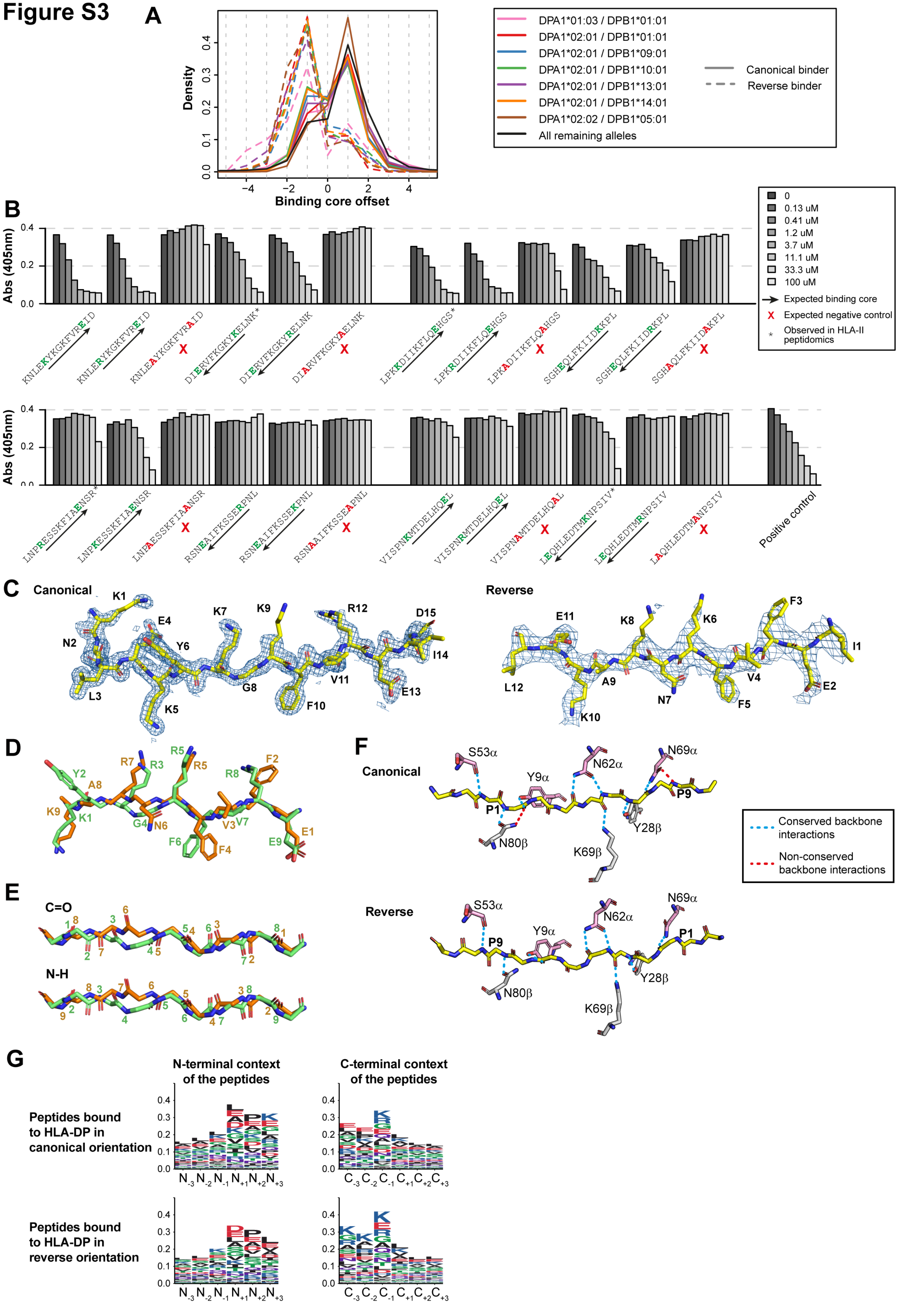

A
Density
Binding core offset
DPA1*01:03 / DPB1*01:01
DPA1*02:01 / DPB1*01:01
DPA1*02:01 / DPB1*09:01
DPA1*02:01 / DPB1*10:01
DPA1*02:01 / DPB1*13:01
DPA1*02:01 / DPB1*14:01
DPA1*02:02 / DPB1*05:01
All remaining alleles
Canonical binder
Reverse binder
B
Abs (405nm)
0
0.13 uM
0.41 uM
1.2 uM
3.7 uM
11.1 uM
33.3 uM
100 uM
Expected binding core
Expected negative control
Observed in HLA-II peptidomics
KNLEKYKGKFVREID
KNLERYKGKFVREID
KNLEAYKGKFVRAID
DIERVFKGKYKELNK*
DIERVFKGKYRELNK
DIARVFKGKYAELNK
LPKKDIIKFLQEHGS*
LPKRDIIKFLQEHGS
LPKADIIKFLQAHGS
SGHEQLFKIIDKKPL
SGHEQLFKIIDRKPL
SGHAQLFKIIDAKPL
LNPRESSKFIAENSR*
LNPKESSKFIAENSR
LNPAESSKFIAANSR
RSNEAIFKSSERPNL
RSNEAIFKSSEKPNL
RSNAAIFKSSEAPNL
VISPNKMTDELHQEL
VISPNRMTDELHQEL
VISPNAMTDELHQAL
LEQHLEDTMKNPSIV*
LEQHLEDTMRNPSIV
LAQHLEDTMANPSIV
Positive control
C
Canonical
Reverse
D
E
C=O
N-H
F
Canonical
Reverse
S53α
Y9α
N62α
N69α
P1
P9
N80β
K69β
Y28β
Conserved backbone interactions
Non-conserved backbone interactions
G
N-terminal context of the peptides
C-terminal context of the peptides
Peptides bound to HLA-DP in canonical orientation
Peptides bound to HLA-DP in reverse orientation

**Figure S3. MHC-II binding specificities reveal a widespread reverse binding mode in MHC-II ligands.**

(A) Distribution of the peptide binding core offsets of canonical versus reverse binding peptides.

(B) Binding competition assays of peptide variants bound to HLA-DPA1*02:01-DPB1*01:01, similar to Figure 3B.

(C) Electron density of the canonical ligand (KNLE**KYKGKFVRE**ID - resolution 1.6 Å) and of the reverse ligand (I**EFVFKNKAK**EL, resolution 2.9Å) obtained in complex with HLA-DPA1*02:01-DPB1*01:01. 2Fo-Fc maps at a contour level of 1 sigma are showed.

(D) Superposition of the binding core of the canonical and reverse ligands, demonstrating clear alignment of the sidechains. For clarity backbone oxygens are not shown.

(E) Superposition of the backbone of the canonical and reverse ligands. The orientation of the backbone N-H and C=O groups are highly conserved, except around P3-P4 of the canonical ligand.

(F) Peptide backbone H-bonds with residues in the MHC-II binding site (top: canonical ligand, bottom: reverse ligand). Conserved H-bonds are shown in blue, non-conserved ones are shown in red.

(G) Motifs of the N- and C-terminal contexts (3 N- and C-terminal residues + 3 residues upstream or downstream of the peptides) of the peptides bound to HLA-DP in the canonical or reverse orientation.

**Figure S4**

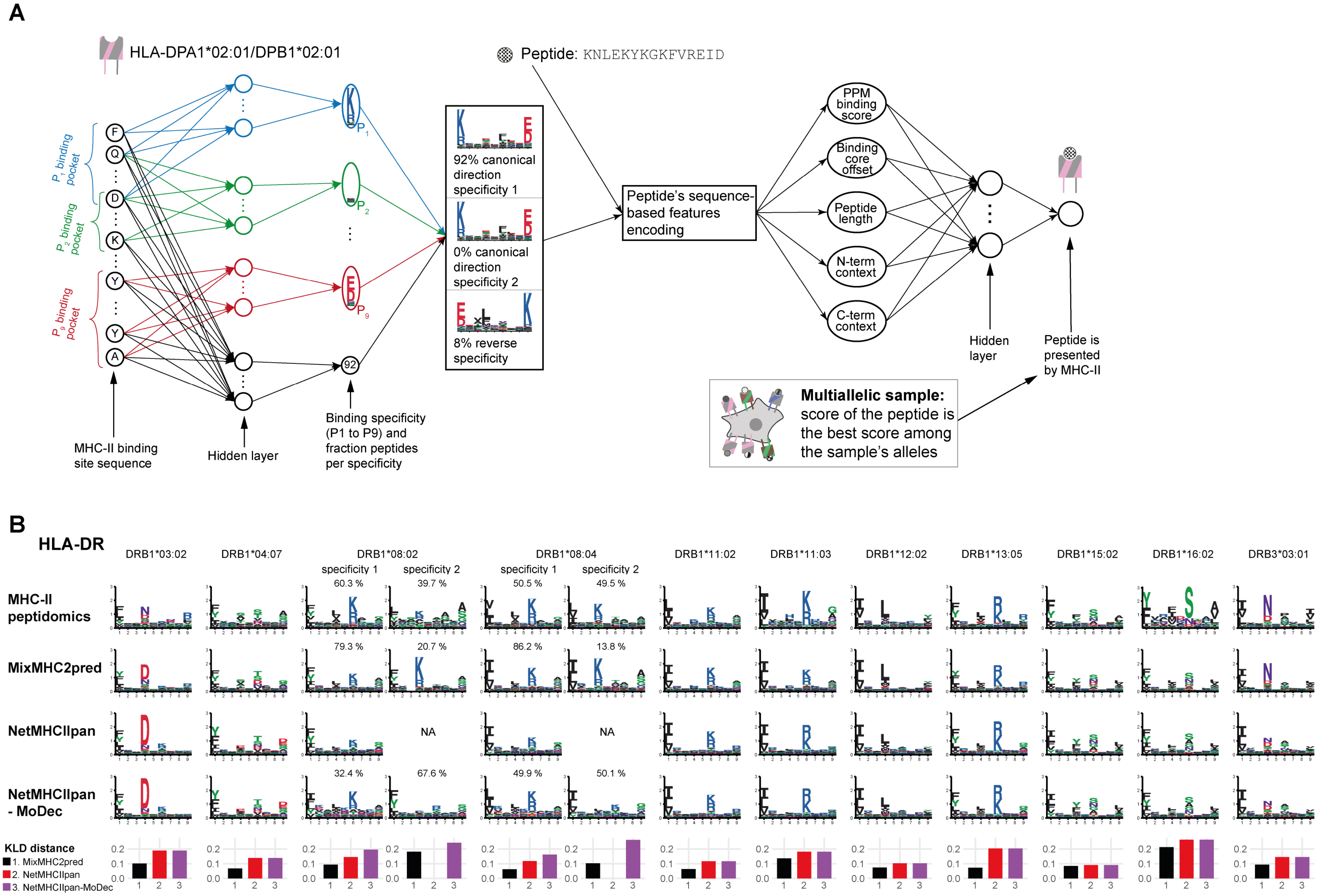

A
HLA-DPA1*02:01/DPB1*02:01
P1 binding pocket
P2 binding pocket
P9 binding pocket
MHC-II binding site sequence
Hidden layer
Binding specificity (P1 to P9) and fraction peptides per specificity
92% canonical direction specificity 1
0% canonical direction specificity 2
8% reverse specificity
Peptide: KNLEKYKGKFVREID
Peptide's sequence-based features encoding
PPM binding score
Binding core offset
Peptide length
N-term context
C-term context
Hidden layer
Peptide is presented by MHC-II
Multiallelic sample: score of the peptide is the best score among the sample's alleles
B
HLA-DR
DRB1*03:02
DRB1*04:07
DRB1*08:02
DRB1*08:04
DRB1*11:02
DRB1*11:03
DRB1*12:02
DRB1*13:05
DRB1*15:02
DRB1*16:02
DRB3*03:01
specificity 1
specificity 2
MHC-II peptidomics
MixMHC2pred
NetMHCIIpan
NetMHCIIpan - MoDec
60.3 %
39.7 %
50.5 %
49.5 %
79.3 %
20.7 %
86.2 %
13.8 %
NA
32.4 %
67.6 %
49.9 %
50.1 %
KLD distance
1. MixMHC2pred
2. NetMHCIIpan
3. NetMHCIIpan-MoDec

**Figure S4. B** (continued)

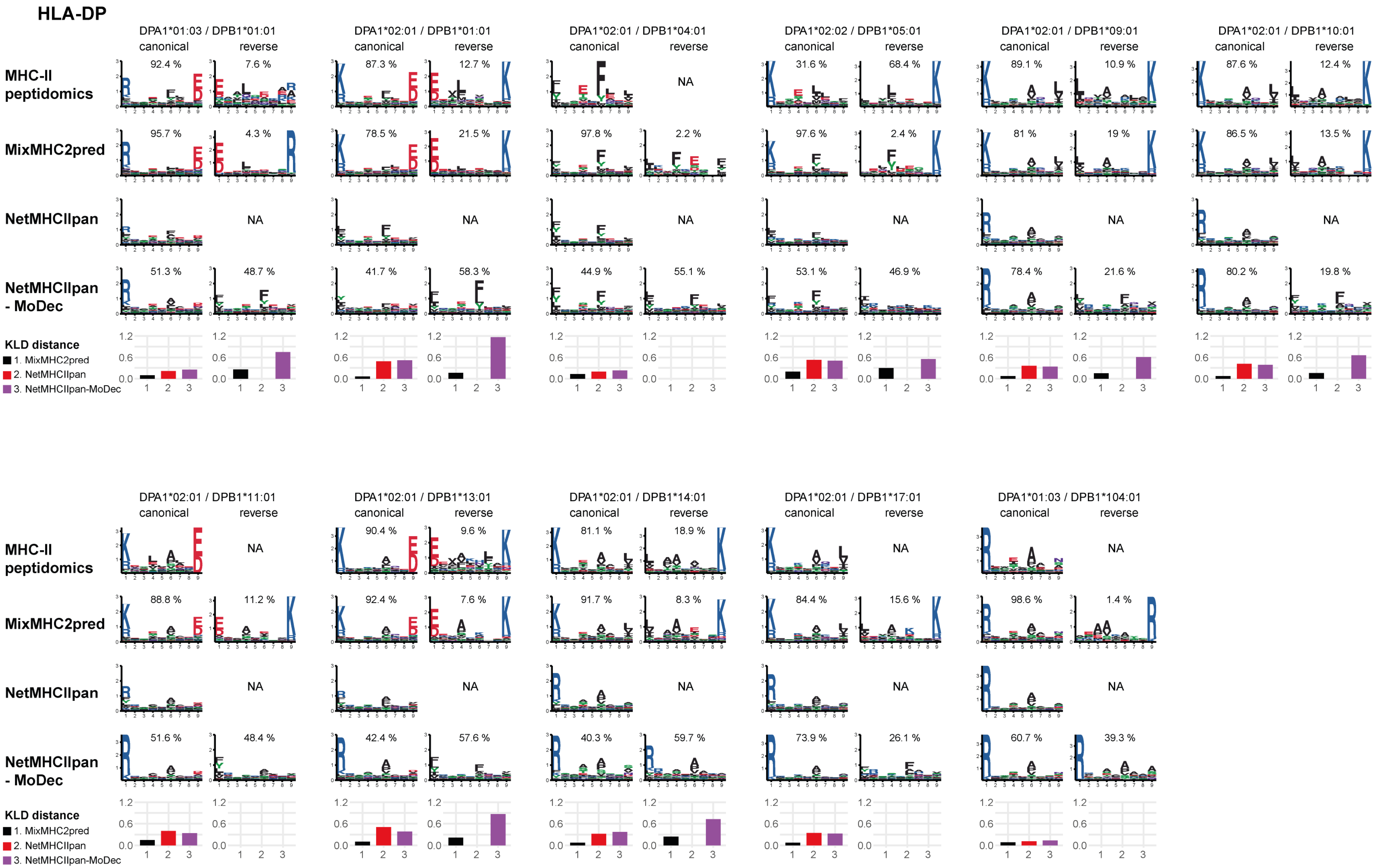

### Figure S4. B (continued)

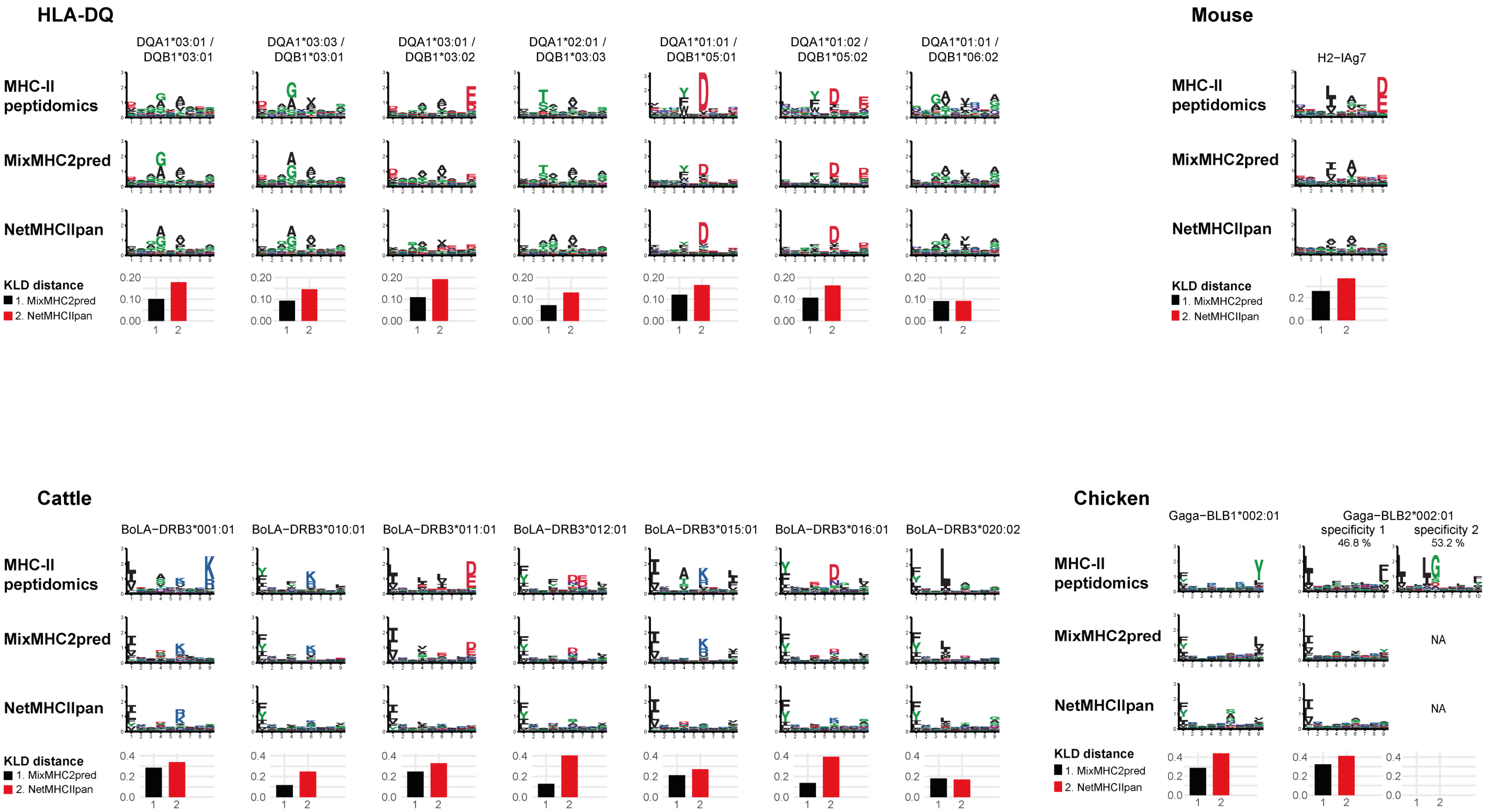

**Figure S4. MHC-II binding specificities can be accurately predicted for alleles without known ligands.**

(A) Description of the pan-allele predictor comprising two consecutive blocks of neural networks. In the first block, which predicts the binding specificity (i.e., PPMs) of an MHC-II allele based on its sequence, multiple independent groups of hidden nodes describe the binding specificity of each peptide binding core position based on the AA residues found in the corresponding binding pocket of the MHC-II allele. A last group of hidden nodes describes the fraction of peptides observed in each binding specificity, based on all AA residues from the MHC-II binding site. The same neural network structure is repeated 3 times to describe the various binding specificities (canonical specificity, second motif in case of bi-specificity, or reverse binding specificity). The PPMs describing the binding specificities of the MHC-II allele are then used to compute binding scores based on the peptide sequence. This score and other features of the peptide are encoded as inputs for the 2$^{nd}$ neural network block. The final output predicts if the given peptide is likely presented by the given MHC-II allele or not. In multiallelic samples, the computations are repeated for each allele of the given sample and the best score determines the final prediction.

(B) Motifs from all alleles absent from NetMHCIIpan-4.0 training predicted in the leave-one-allele-out cross-validation. When multiple specificities are present, the fraction of peptides observed and predicted per motif is indicated above each motif (for NetMHCIIpan, two rows are showed: 1$^{st}$ corresponds to results from this predictor directly – without multiple specificity as not indicated in its returned results; 2$^{nd}$ row is when applying MoDec on the best predicted cores of NetMHCIIpan, searching for 2 motifs). The average Kullback-Leibler divergence (KLD) per peptide binding core position between the specificities observed in MHC-II peptidomics and predicted ones is showed below each allele.

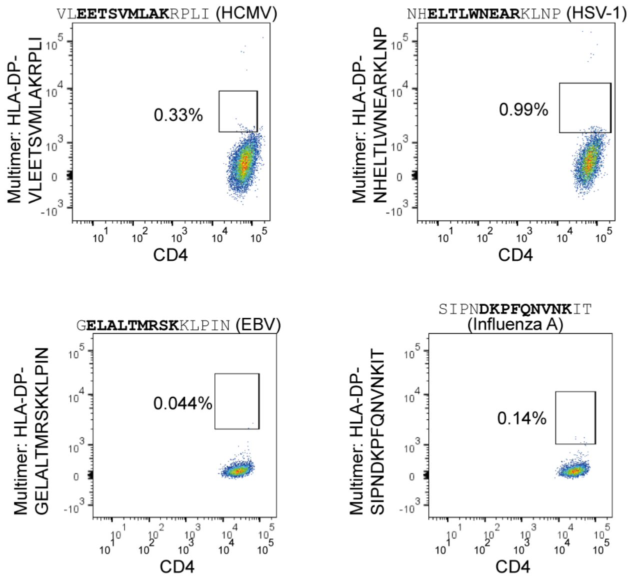


**Figure S5. Multiple specificities reveal reverse binding CD4+ T-cell epitopes.** Negative controls based on irrelevant HLA-matched donors, relating to the peptide-MHC-II multimers of the reverse binding epitopes from Figure 6C.

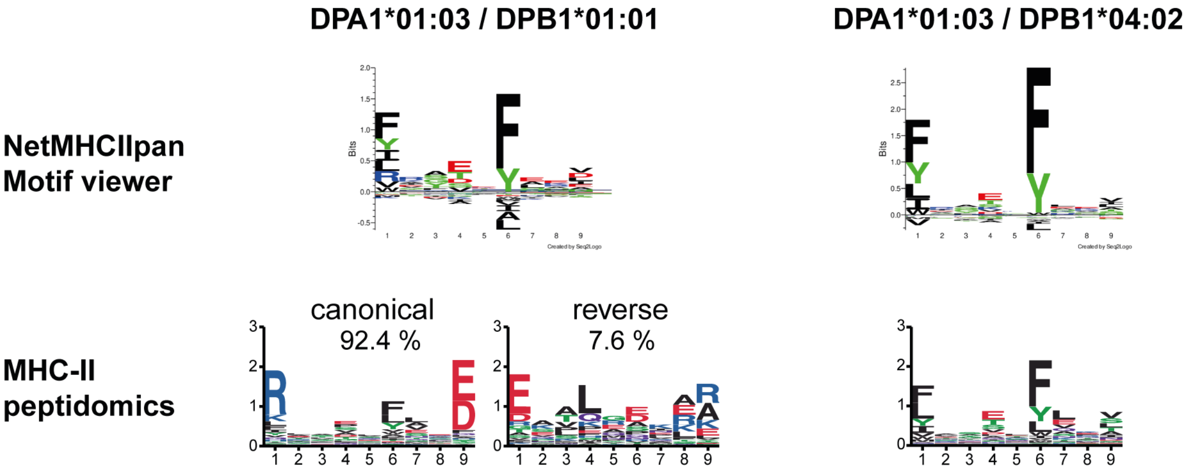


**Figure S6. Supertypes do not necessarily reflect biological properties of MHC-II alleles.** HLA-DPB1*01:01 and HLA-DPB1*04:02 are found in the same supertype defined in (Greenbaum et al., 2011) and their binding specificity is predicted to be similar based also on NetMHCIIpan Motif viewer (top motifs; https://services.healthtech.dtu.dk/service.php?NetMHCIIpan-4.0) (Reynisson et al., 2020). This is refuted by our large MHC-II peptidomics dataset (bottom motifs) showing large deviations between the binding specificities of the two alleles, especially at their P1 and P9 peptide binding cores.

**Table S1. Information about the different cell lines and tissue samples used in the HLA-II peptidomics experiments from this study and curated from public MHC-II datasets.** (Table is available as an Excel file).

**Table S2. MHC-II binding specificities reflect biochemical properties of the MHC-II binding pockets.** (Table is available as an Excel file).

(A-C) Sequences of the amino acids in the HLA-DR (A), HLA-DP (B) and HLA-DQ (C) binding pockets and corresponding peptide binding specificities.

(D) Discussion about the observed specificities.

(E) Calculated change in FoldX energy score for several variants of DRB1*08:01 and DRB1*13:01 ligands.

**Table S3. MixMHC2pred-2.0 unravels new reverse binding CD4+ T-cell epitopes.** (Table is available as an Excel file).

(A) Reverse binding candidates tested for immunogenicity, the observed immunogenicity response in two donors (based on TNFα and IFNγ production) and their score with various predictors.

(B) TCR sequencing of the sorted $CD4^+$ T cells responding to the observed epitopes.

**Table S4. Data collection and refinement statistics of the crystal structures obtained in this study.**

| **Structure** | **HLA-DP - peptide (canonical orientation)** | **HLA-DP - peptide (reverse orientation)** |
|---|---|---|
| **PDB ID** | **XXXX** | **YYYY** |
| ***Data Collection*** | | |
| Resolution (Å) | 48.11-1.62 (1.71-1.62)* | 45.39-2.90 (3.07-2.90)* |
| Space group | $P2_12_12_1$ | $P6_122$ |
| Cell dimensions | | |
| a, b, c (Å) | 54.00, 82.68, 105.98 | 83.98, 83.98, 348.59 |
| a, b, g (°) | 90, 90, 90 | 90, 90, 120 |
| Unique reflections | 116997 | 16932 |
| $R_{meas}$ | 0.084 (1.97) | 0.192 (1.095) |

| | | |
|---|---|---|
| <I/s(I)> | 12.43 (0.83) | 11.55 (1.82) |
| Data completeness (%) | 99.7 (98.5) | 97.9 (99.7) |
| $CC_{1/2}$ | 0.999 (0.404) | 0.996 (0.816) |
| Redundancy | 6.7 (6.5) | 10.6 (10.6) |
| ***Refinement Statistics*** | | |
| $R_{work}$ | 0.205 | 0.231 |
| $R_{free}$ | 0.231 | 0.272 |
| RMSD from ideality | | |
| Bond lengths (Å) | 0.01 | 0.002 |
| Bond angles (°) | 1.046 | 0.565 |
| Ramchandran statistics (%) | | |
| Favored | 99.2 | 96.3 |
| Allowed | 0.5 | 3.5 |
| Outlier | 0.3 | 0.3 |

*Values in parenthesis are for the highest resolution shell.

**Data S1. List of MHC-II ligands obtained:**

(A) from public MHC-II datasets.

(B) in the new HLA-DR, HLA-DP, HLA-DQ and remaining pan-HLA-II peptidomics data from our study.

(Data S1 is available as an Excel file).
